# Epigenomic characterization of latent HIV infection identifies latency regulating transcription factors

**DOI:** 10.1101/2020.07.24.220012

**Authors:** Stuart R Jefferys, Sam Burgos, Jackson J Peterson, Sara R Selitsky, Anne-Marie Turner, Lindsey I James, David M Margolis, Joel Parker, Edward P Browne

## Abstract

Transcriptional silencing of HIV generates a reservoir of latently infected cells, but the mechanisms that lead to this outcome are not well understood. We characterized a primary cell model of HIV latency, and observed that latency is a stable, heritable viral state that is rapidly reestablished after stimulation. Using Assay of Transposon-Accessible Chromatin sequencing (ATACseq) we found that latently infected cells exhibit reduced proviral accessibility, elevated activity of Forkead and Kruppel-like factor transcription factors (TFs), and reduced activity of AP-1, RUNX and GATA TFs. Latency reversing agents caused distinct patterns of chromatin reopening across the provirus. Furthermore, depletion of a chromatin domain insulator, CTCF inhibited HIV latency, identifying this factor as playing a key role in the initiation or enforcement of latency. These data indicate that HIV latency develops preferentially in cells with a distinct pattern of TF activity that promotes a closed proviral structure and inhibits viral gene expression.

## Introduction

HIV infection continues to be a major global health problem, with 37 million infected individuals and approximately one million deaths per year (Yoshimura, 2017). HIV infection can be potently controlled by combination antiretroviral therapy (ART), with treatment suppressing viral loads to undetectable levels and allowing HIV-infected patients to live lives of roughly normal lifespan. Nevertheless, persistent health complications in treated patients, as well as side effects of ART, make the development of a cure for HIV a high priority (Lederman et al., 2011, 2013; Lichtfuss et al., 2011; Serrano-Villar et al., 2014). Moreover, rapid rebound of viremia after treatment interruption demonstrates the persistence of HIV-infected cells, even after long-term ART. Indeed, longitudinal studies to estimate the half-life of this reservoir indicate that greater than 72 years of treatment would be necessary for elimination of all infected cells (Crooks et al., 2015; Siliciano et al., 2003). The mechanisms of HIV persistence are not completely understood, but likely involve several factors. First, HIV is able to establish a latent infection, characterized by the presence of a transcriptionally silent provirus, in a subset of host cells (Chun et al., 1997; Finzi et al., 1997). Sporadic reactivation of these cells may occur continuously, and leads to viral rebound if treatment is interrupted (Pinkevych et al., 2015). The precise location of this latently-infected reservoir is debated, but is likely distributed across a number of CD4 T cell subsets, including long-lived memory cells, thereby explaining the long persistence of infection (Baxter et al., 2016; Lee and Lichterfeld, 2016; Soriano-Sarabia et al., 2014). Second, HIV-infected cells can undergo homeostatic expansion *in vivo* during ART, further replenishing the pool of infected cells (Chomont et al., 2009; Maldarelli et al., 2014; Reeves et al., 2018). Understanding the molecular mechanisms of how latency is established and maintained will be critically important to developing strategies to prevent or eliminate the latent reservoir. Certain cell types, such as resting memory CD4 T cells, are a suboptimal environment for HIV transcription, due to limiting availability of transcription factors required for HIV gene expression, including NF-κB, AP-1 and P-TEFb (Dahabieh et al., 2014; Kim et al., 2011; Nabel and Baltimore, 1987; Tyagi et al., 2010). Stochastic variation in the levels of the viral Tat protein during infection may also contribute to latency (Razooky et al., 2015). Furthermore, covalent modification of provirus–associated histones by histone-modifying enzymes such as histone deacetylases (HDACs) or histone methyl transferases (HMTs), such as the PRC2 or HUSH complexes, can have a repressive effective on viral transcription (Archin et al., 2012; Barton et al., 2014; Chougui and Margottin-Goguet, 2019; Friedman et al., 2011; Tripathy et al., 2015), and their role in the maintenance of latency is strongly supported by evidence of latency reversal in vivo (Mzingwane and Tiemessen, 2017).

We have previously established, using a primary CD4 T cell model of HIV latency, that latency can occur in diverse host cell environments, but occurs preferentially in cells that express markers of quiescent central memory T cells (Tcm), and that exhibit high proliferative potential (Bradley et al., 2018). These results indicate that the establishment of latency is influenced by the intrinsic biological program of the host cell. However, the mechanistic details of how specific host cell environments or phenotypes impact the initiation or maintenance of latency are unknown. To further investigate this observation, we sought to characterize primary CD4 cells in which latency has become established by defining chromatin-based characteristics of these cells. The results revealed an association of the latency phenotype with a distinct pattern of chromatin accessibility and reduced accessibility of the HIV genome. Furthermore, we identify a set of cellular transcription factors with differentially accessibly binding sites in latently infected cells, which may play a role in influencing the course of HIV transcriptional silencing and the entry into or maintenance of the latent state. In particular, we investigate and confirm the role of CTCF during HIV latency, implicating this protein as novel latency-regulating factor.

## Results

### HIV-infected cells enter a stable, heritable state of latency in a cell culture model

We have previously established a cell culture model of HIV latency (**Figure 1A**). In this model, primary CD4 T cells are activated through TCR stimulation, then infected with a reporter HIV strain that encodes a destabilized eGFP gene (herein referred to as HIV-GFP) (Yang et al., 2009). Actively infected (GFP+) cells are purified by flow sorting at 2dpi, and then cultured for up to 12 weeks. During this time period, viral gene expression progressively diminishes, and a subset of the cells become GFP- (latently infected), while the remaining cells exhibit variegated levels of viral gene expression. To examine the stability of the viral gene expression phenotype, we resorted this mixed culture at 12wpi into GFP+ (actively infected) and GFP- (latently infected) (**Figure 1B**) and cultured these cells independently for six additional days. Notably, the infected cells retained their viral gene expression level, that is GFP+ cells remained GFP+, and GFP-cells remained GFP-, for this time period indicating that the level of viral gene expression is a relatively stable property of the infected cells (**Figure 1C**). To look at the effect of cellular stimulation on viral gene expression in latently infected cells, we isolated the latently infected (GFP-) cells by flow sorting, and then restimulated them through their TCR for 3 days, monitoring GFP expression over time (**Figure 1D**). At the time of stimulus removal, the latently infected cells had potently upregulated GFP expression and a large fraction of the cells (90%) had become GFP+. Interestingly, after removal of the TCR stimulus, the majority of reactivated cells then rapidly lost viral gene expression and returned to latency (GFP-) within 14 days. A residual population of around 20% of the reactivated cells retained a low level of GFP expression after this single round of reactivation. To further investigate this finding, we stimulated these cells three additional times over a total period of 48 days (**Figure 1E**). Similar to the initial restimulation, after each subsequent stimulation the latently infected cells rapidly upregulated GFP expression, then returned to latency with highly consistent kinetics after stimulus removal. Specifically, the cells returned to a largely GFP-phenotype by 7d post stimulation. Notably, during this period of the time the cells also underwent substantial cell division and cell numbers increased by ~170 fold (**Figure 1F**). This finding suggests that latency is a stable, intrinsic property of infected cells that is rapidly re-established after viral reactivation, and can be transmitted to daughter cells during cell division. This conclusion is consistent with previous studies showing rapid reestablishment of latency after stimulation of infected cells with histone deacetylase inhibitors (Shan et al., 2014).

**Figure 1:**
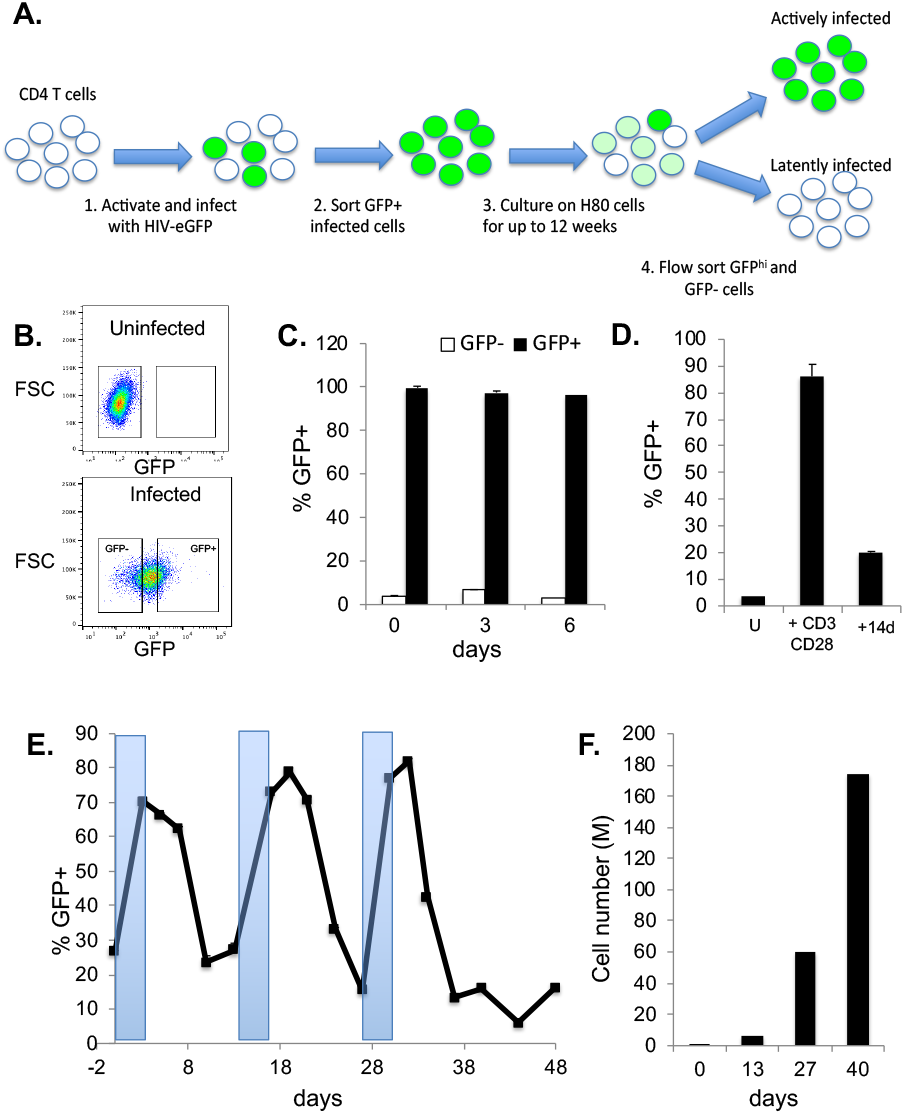
HIV infected CD4 T cells enter a stable, heritable state of viral latency in vitro. **A.** Schematic overview of primary CD4 T cell HIV latency model and **(B)** flow cytometry plot showing a representative gating of actively infected (GFP+) and latently infected (GFP-) cells. **C.** GFP+ and GFP-cells were flow sorted from the HIV-GFP infected population and cultured for 6 days post sorting. Viral gene expression was measured at days 0, 3, and 6 post sorting, and the percentage of cells in the GFP+ gate was measured. Each bar represents the average of three independent replicates. **D.** Sorted infected GFP-cells at 12wpi were reactivated using αCD3/CD28 beads. At 24h, unstimulated cells (U) and stimulated cells (+αCD3/CD28) were analyzed by flow cytometry and the percentage of GFP+ cells analyzed. The reactivated cells were then removed from the activating beads and cultured for an additional 14 days (+14d), then analyzed again by flow cytometry. **E.** Sorted GFP-cells from an infected population were serially stimulated through their TCR (+αCD3/CD28) and GFP expression was monitored over time by flow cytometry. Shaded areas indicate times during which αCD3/CD28 beads were added to the culture to stimulate the cells. Each datapoint represents the average of two independent replicates. **F**. Cell numbers were counted at selected timepoints during the serial stimulation shown in **E** and total cell numbers are shown in millions of cells (M).

### Latently infected cells exhibit differentially accessible chromatin

The stable, heritable character of HIV latency in this model system lead us to hypothesize that epigenomic or chromatin-based changes in infected cells were associated with reversible but recurring viral silencing. Gene activity can be regulated by changes in the structure and accessibility of chromatin. These changes can be mediated by chromatin remodeling complexes that are recruited to sites of active transcription by transcription factors (TFs) and result in the removal or repositioning of nucleosomes near transcription start sites (TSS) (He et al., 2002). Furthermore, enzymes that add or remove covalent modifications to histones tails can create a histone “code” that affects the structure of the chromatin and creates docking sites for additional regulators (Strahl and Allis, 2000). Changes in chromatin structure and accessibility can then allow greater access to the promoter by core transcriptional regulators including RNA polymerase, thereby facilitating transcription.

To examine whether the establishment of latency in our model system was associated with specific changes in chromatin accessibility, both within the HIV genome and within the host cell genome, we performed Assay of Transposon-Accessible Chromatin sequencing (ATACseq) on infected cells. ATACseq uses a hyperactive Tn5 transposon to probe and identify regions of open chromatin (Buenrostro et al., 2015a, 2015b). To compare cells with latent infection to those with active viral transcription, we sorted HIV-GFP infected cells at 12 weeks after infection based on viral gene expression into GFP+ or GFP-populations, representing cells from the upper and lower 25% of GFP expression intensity respectively, and generated ATACseq libraries from the sorted cells. To ensure biological reproducibility, we analyzed infected cells derived from two independent donors. These data were then aligned to a combined reference genome including the human genome and HIV. Quality control analyses showed nucleosomal laddering of fragment sizes and enrichment of reads at transcription start sites (TSS), indicating high quality data (**Figure 2A, 2B**). Replicate analysis of cells derived from the same donor clustered closely together by principal component analysis (PCA), indicating high reproducibility (**Figure S1**).

**Figure 2:**
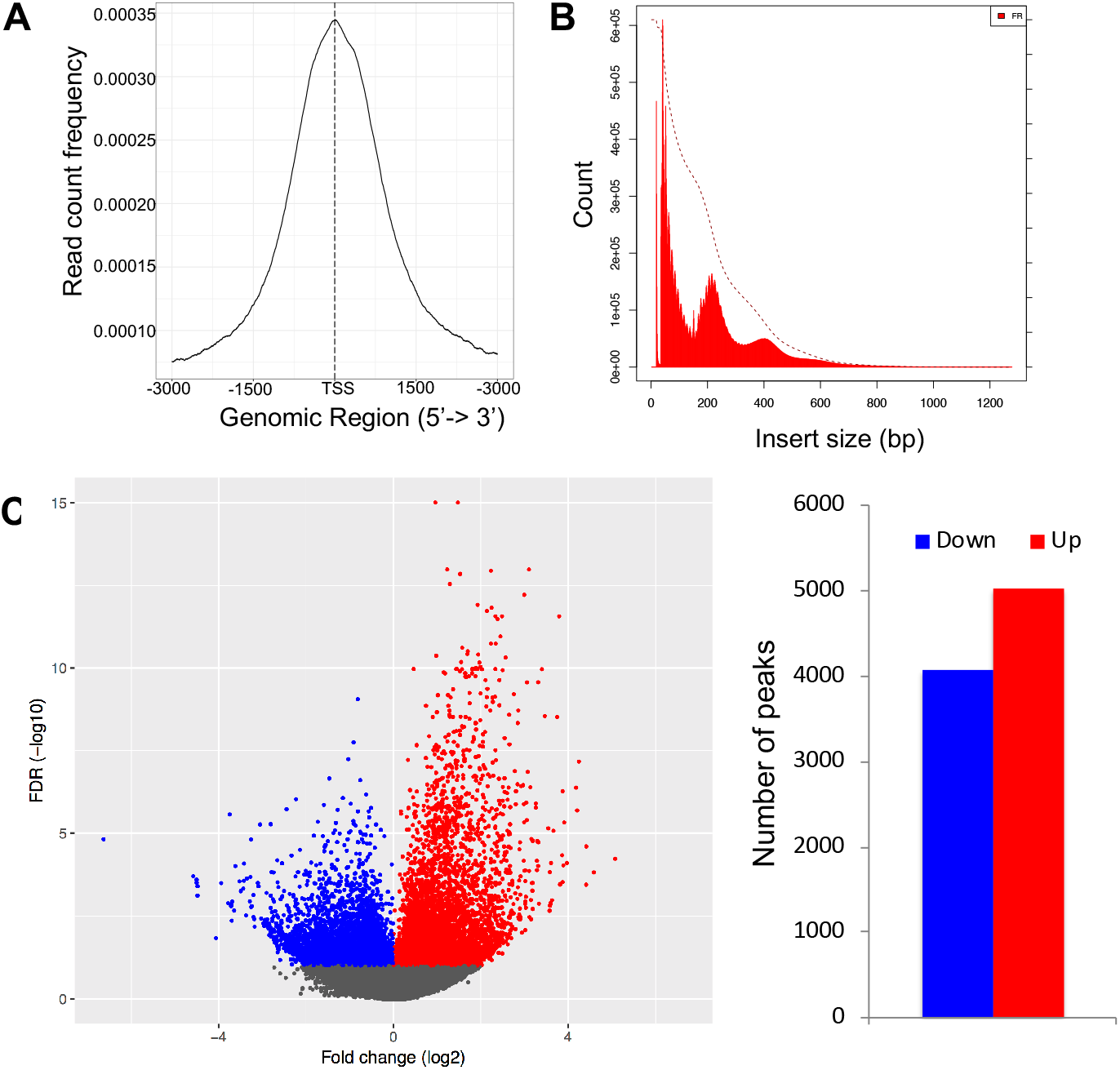
Latently infected cells are epigenomically distinct from actively infected cells. HIV-GFP infected cells at 12 wpi were flow sorted into GFP- and GFP+ populations and analyzed by ATACseq. **A.** Alignment of the reads to the human genome demonstrates enrichment of reads near transcriptional start sites (TSS). **B.** Size distribution of library fragment sizes showing nucleosomal laddering. **C**. Differentially accessible regions of the cellular genome were identified and fold change versus false discovery rate (FDR) displayed as a volcano plot. Peaks that are significantly (FDR <0.1) more open in GFP+ cells (“up”) are indicated in red, while peaks that are significantly more open in latently infected cells (“down”) are indicated in blue. The data represents an aggregate of infected cells from two independent donors.

Next, we compared accessibility of chromatin from the GFP+ and GFP-infected cells in greater detail by calculating differential accessibility for each of the accessibility peaks present in the samples. These data showed that GFP- and GFP+ cells have distinct chromatin accessibility patterns, with several thousand differentially open peaks between latently infected and actively infected cells (**Figure 2C**). Specifically, using a false discovery rate (FDR) cutoff value of 0.05, we observed 5021 peaks that were significantly more open in GFP+ cells, and 4068 peaks that were more open in GFP-cells, out of 277992 overall peaks detected across all samples (**Figure 2C**). These data thus indicate that viral latency in this system is correlated with a specific pattern of overall genomic accessibility.

### The HIV genome exhibits reduced accessibility in latently infected cells

Next we aligned the ATACseq reads to the HIV genome to determine whether HIV latency is associated with differential accessibility of the provirus.. For both GFP+ and GFP-infected cells the HIV genome exhibits major accessibility peaks surrounding the 5’ transcription start site (TSS) at nucleotide (nt) 455, and within the 3’ LTR (**Figure 3A**). Downstream of the TSS peak, after nt 800, the virus genome becomes significantly less accessible, consistent with the hypothesis that chromatin structural barriers proximal to TSS impede processive viral transcription (Battivelli et al., 2018; Rafati et al., 2011). We also observed an overall increasing gradient of openness across the 3’ half of virus in actively infected cells, leading to the large peak in the 3’LTR region. (**Figure 3A**). Overall we observed significant reduced accessibility of the HIV provirus in the latently infected cells relative to the actively infected cells (2.45 fold, FDR=2.87×10^−13^).To identify specific regions within the HIV genome with differential accessibility, we subdivided the HIV genome into 73bp bins and individually calculated the fold change and false discovery rate for each bin (**Figure 3B**). This analysis showed significantly reduced accessibility across the viral genome in latency, but a particularly large difference towards the 3’ end of the provirus. This finding is consistent with the hypothesis that physical barriers limiting the accessibility of the provirus to factors that regulate transcription likely contribute to latency.

**Figure 3:**
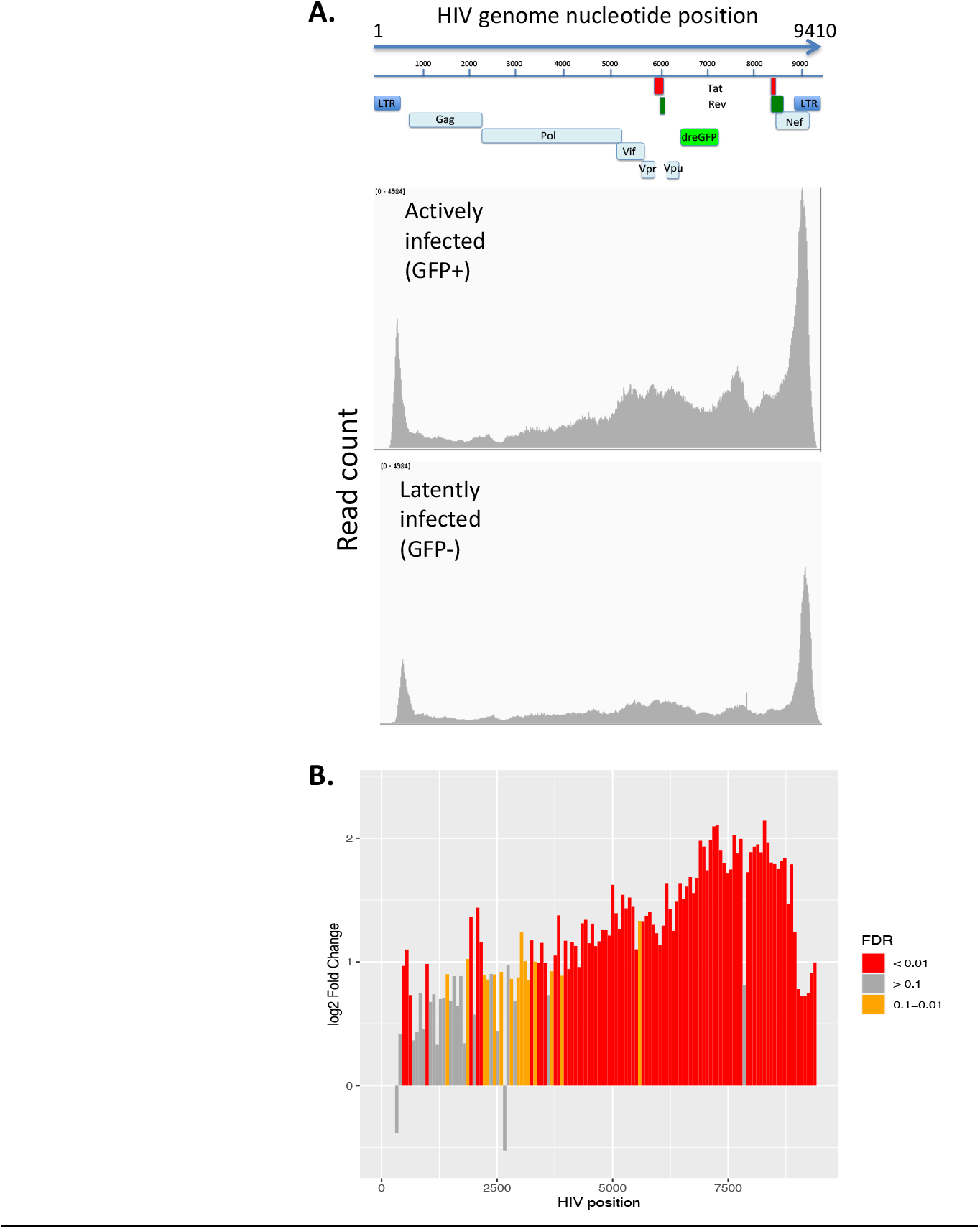
The HIV genome has reduced accessibility in latently infected cells. **A.** HIV-GFP infected cells at 12wpi were flow sorted into GFP- and GFP+ populations and analyzed by ATACseq. ATACseq reads from the cells were aligned to an HIV reference genome to examine accessibility of the provirus. The intensity of read coverage was graphed across the HIV genome for GFP+ (upper panel) and GFP- (lower panel) cells. Graph shows an aggregate analysis of two donors. **B.** The change in accessibility across the HIV genome during active infection vs latency is shown by determining the fold change between GFP- and GFP+ cells within 73bp bins tiled across HIV. Positive fold change indicates greater accessibility in GFP+ cells. The false discovery rate (FDR) for each 73bp bin is illustrated by the color of each bar: red=<0.01, orange=0.1 to 0.01, grey= >0.1.

### Latency reversal is associated with changes in accessibility in the host cell and viral genomes

We next examined the impact of latency reversing agents (LRAs) on latently infected cells. We stimulated HIV-GFP infected cells at 12wpi with three different LRAs, each with a different mechanism of action – vorinostat (histone deacetylase inhibitor), AZD5582 (SMAC mimetic), and prostratin (PKC agonist) for 24 hr. We have previously examined the transcriptomic impact of these three LRAs in primary CD4 T cells (Nixon et al., 2020). At 24h, we measured the response of viral gene expression to the LRAs by flow cytometry. Vorinostat and AZD5582 caused modest increases (~5-10%) in the percent GFP+ cells in the culture, indicating weak latency reversal. By contrast, prostratin caused a more substantial increase in viral gene expression (~25%) (**Figure 4A**).

**Figure 4:**
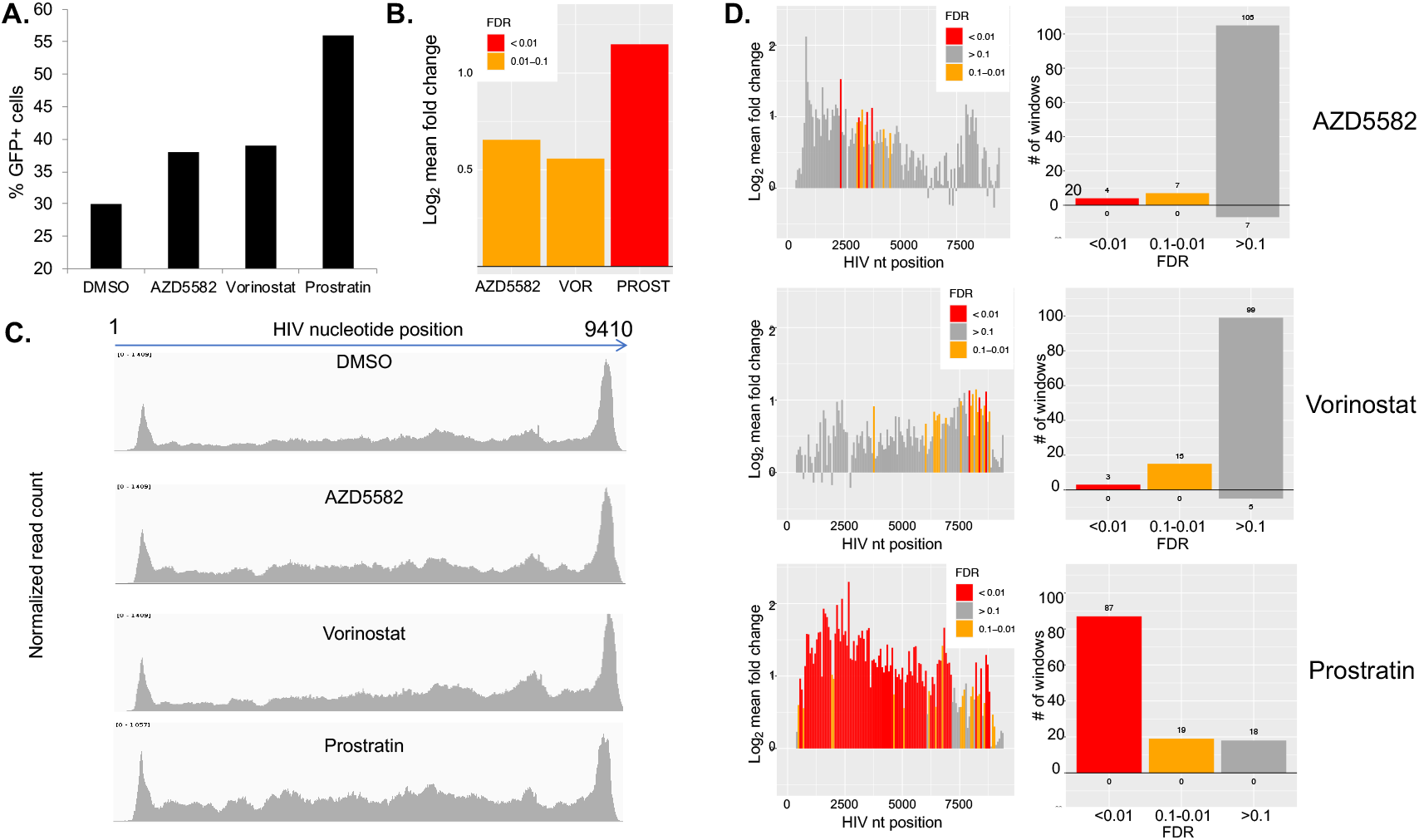
Latency reversal is associated re-opening of the viral genome. HIV-GFP infected CD4 cells at 12wpi were stimulated with three different latency reversing agents – vorinostat (250nM), AZD5582 (250nM), prostratin (250nM) or DMSO (0.1%) vehicle for 24hrs **A.** GFP expression in the culture was analyzed by flow cytometry and the percent GFP+ was plotted. **B.** Stimulated cells were then analyzed by ATACseq. Reads were aligned to the HIV genome and the fold change of reads mapping to HIV in each of the LRA-stimulated conditions are shown. VOR=vorinostat, PROST=prostratin. **C.** The read coverage across the HIV genome for the four conditions (DMSO control and the LRA conditions) are shown. **D**. Fold change in accessibility after LRA stimulation is shown by examination of 73bp bins across the genome (Left panels). Individual bars are color coded by the false discover rate (FDR): red=<0.01, orange=0.1 to 0.01, grey= >0.1. The total number of differentially accessible 73bp bins across the genome in each FDR category are shown in the right panel. Data represent an aggregate of four independent replicate experiments.

We then performed ATACseq on the LRA stimulated cells, as well as on cells that had been exposed to vehicle (DMSO 0.1%) only. We first examined the impact of LRA stimulation on accessibility of the HIV genome. Notably, all three LRAs caused an increase in fraction of reads that mapped to the HIV genome, although trend was more pronounced for prostratin (**Figure 4B, 4C**). Specifically, vorinostat and AZD5582 caused 1.47 fold (FDR= 1.49×10^−2^) and 1.57 fold (FDR=8.81×10^−2^) increases in proviral accessibility respectively, while prostratin caused a 2.22 fold increase, (FDR=1.871×10^−8^). This observation suggests that latency reversal is associated with reopening of the closed latent provirus, and that the potency of LRAs is related to the magnitude of proviral reopening that they induce. To examine changes in viral accessibility after LRA stimulation in more detail, we subdivided the HIV genome into 73bp bins and individually calculated the fold change and false discovery rate for each bin (**Figure 4D**). Interestingly, each LRA induced a distinct pattern of changes to proviral accessibility. Prostratin and AZD5582 caused significantly increased accessibility mainly towards the 5’ end of the virus, while vorinostat-induced reopening occurred mainly towards the 3’ end of the virus. (**Figure 4D**). The biological basis for the differential behavior of the LRAs in this analysis is unknown, but suggests the existence of previously unappreciated mechanisms of interaction between each agent and the provirus.

By alignment of data from the LRA stimulated cells to the human genome, we were able to identify regions of cellular chromatin for which accessibility was modulated by each agent. PCA analysis of accessibility peak data from the four conditions showed that replicate samples stimulated with a given LRA tended to cluster together (**Figure S2**). Prostratin and AZD5582 stimulated cells formed distinct clusters separated from each other and vorinostat by PC2. Vorinostat and control vehicle (0.1% DMSO) stimulated cells overlapped in the plot indicating a more limited impact of vorinostat on the accessibility of the host cells. Prostratin had the most dramatic effect on the cells, consistent with its known role as a potent modulator of cellular transcription. Analysis of individual peak dynamics by volcano plot also revealed that each LRA induced a distinct pattern of changes to the accessible chromatin of infected cells (**Figure 5**). Prostratin had the most pronounced impact, and caused numerous accessibility peaks to open and to close. Specifically, 33723 peaks were more open, while 22458 peaks more closed after 24h prostratin stimulation. By contrast, vorinostat caused much more modest changes to host cell chromatin accessibility, with 2326 peaks more open and 3323 peaks more closed. Interestingly, the effects of AZD5582 on chromatin accessibility were highly asymmetric, with 11362 peaks more open and 4945 peaks more closed. This observation is consistent with our previous observation of the asymmetric effect of this compound on transcript levels (Nixon et al., 2020).

**Figure 5:**
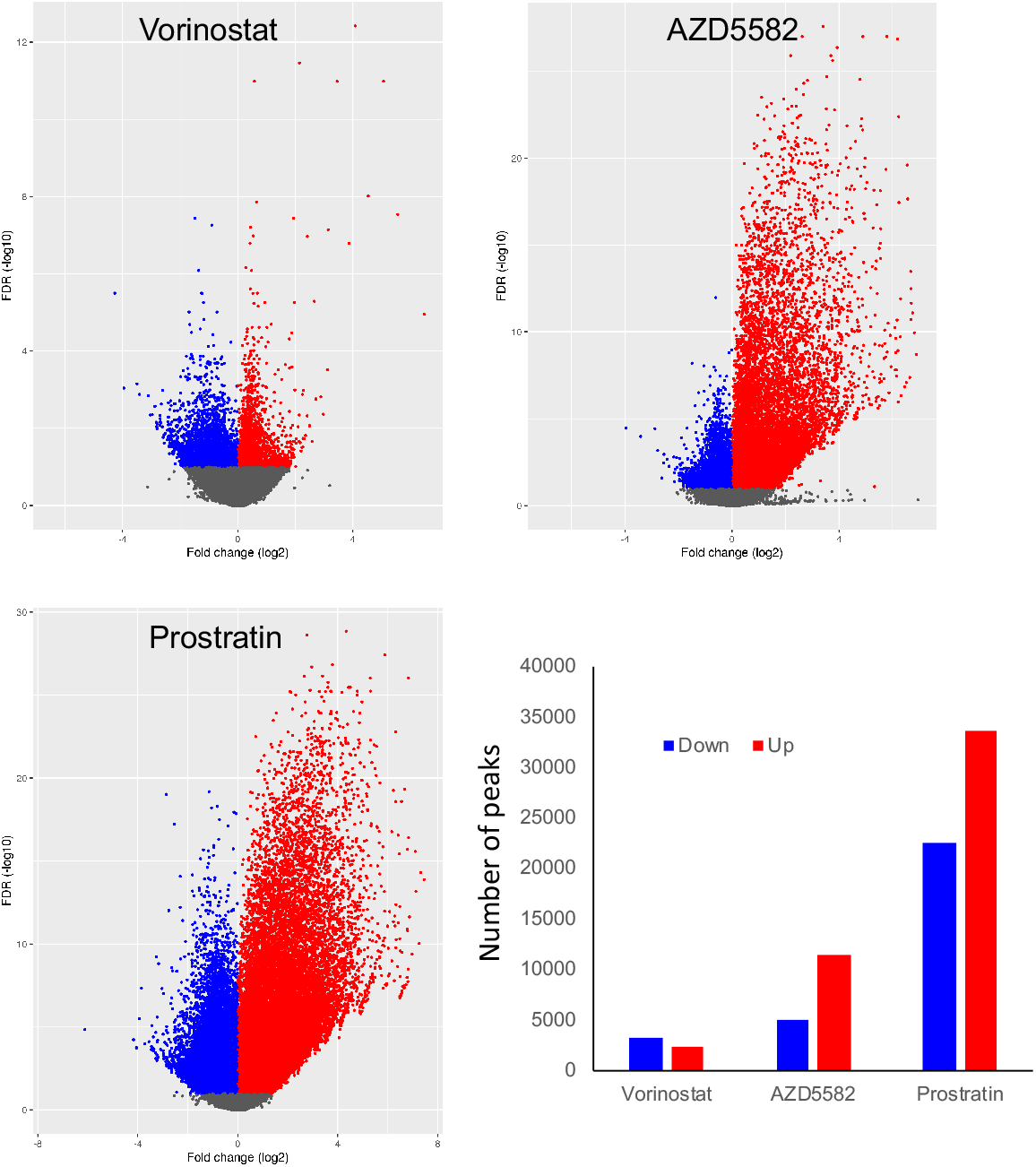
LRAs induce changes to chromatin accessibility in infected cells. ATACseq read from HIV-GFP infected cells at 12wpi that had been stimulated with Vorinostat (250nM), AZD5582 (250nM), Prostratin (250nM) or DMSO vehicle for 24hrs were aligned to the human genome. Differentially accessible regions were identified and fold changes vs FDR plotted as a volcano plot. Data represent an aggregate analysis of four independent replicate experiments. Fold changes versus FDR are displayed as a volcano plot. Peaks that are significantly (FDR < 0.1) more open after LRA stimulation are indicated in red (“up”), while peaks that are significantly more closed are indicated in blue (“down”).

### Differential accessibility of specific transcription factor binding sites is associated with latency and latency reversal

To further investigate the regulation of HIV latency, we next examined the enrichment of specific transcription factor (TF) binding sites in the differentially accessible chromatin peaks identified by analysis of the ATACseq data. Since active transcription factors typically promote local opening of chromatin surrounding their binding sites, this information can be used to infer the activity of specific TFs in cells of interest. To investigate this, we analyzed the set of variable peaks using a motif enrichment method, HOMER (Boeva, 2016). This method analyzes defined DNA regions for statistical enrichment of known TF binding site sequences, relative to a background reference sequence. First, we compared the TF site enrichment in the DNA sequences of differential accessible peaks between actively infected (GFP+) and latently infected (GFP-) cells, using the non-differentially accessible peaks as a background baseline (FDR>0.2). From this, we derived a list of transcription factors with binding sites that were enriched in the peaks that were differentially open between latently infected and actively infected cells (**Table 1, Table S1, Table S2)**. We identified 406 potential TF binding sequences that were preferentially enriched (P <0.05) in GFP+ cells and 252 sequences that were enriched in GFP-cells. Notably, many of the most highly enriched motifs were variants of consensus binding sites for members of large TF families. Since the binding sites of the individual members in these families are often very similar, precise disambiguation of which specific TFs are active in these cells is difficult to achieve from this data alone. Nevertheless, from examination of the 20 most enriched binding site motifs for each condition, several clear trends were noticeable from the data. Actively infected (GFP+) cells displayed strongly elevated activity for three TF families –AP-1 related TFs, Runt-related (RUNX) TFs, and GATA TFs. Latently infected (GFP-) cells, by contrast, displayed elevated activity of Kruppel-like TFs (KLFs), and Forkhead (FOX) TFs. Notably, several members of these TF families have known roles in regulating HIV gene expression indicating that the results are biologically valid. In particular, AP-1 has well characterized binding sites within the HIV promoter, and has been demonstrated to promote HIV transcription (Duverger et al., 2013; Roebuck et al., 1996; Yang et al., 1999). These results are also consistent with our previous scRNAseq findings in this model, which found elevated and FOXP1 expression in latently infected cells, and elevated GATA3 expression in actively infected cells (Bradley et al., 2018).

**Table 1:**
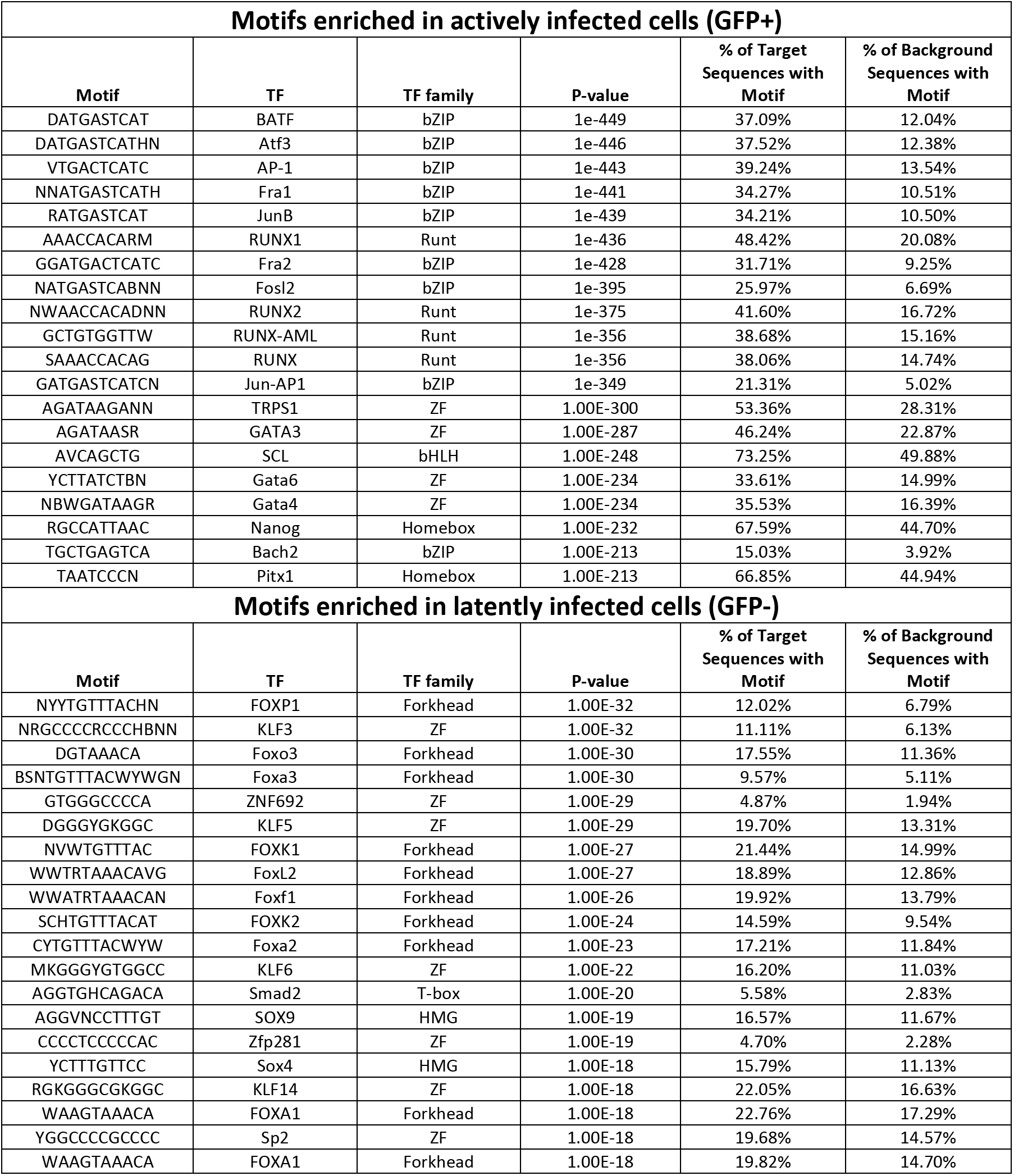
TF motifs enriched in accessible chromatin of GFP+ vs GFP-cells. HOMER motif enrichment analysis of latently infected (GFP-) cells versus actively infected (GFP+) cells, using the total set of open peak sequences in CD4 T cells as a reference baseline The 20 most enriched sequences in each population relative to the other, and the putative TFs that bind these sequences are displayed. Full list found in Table S2.

**Table 2:**
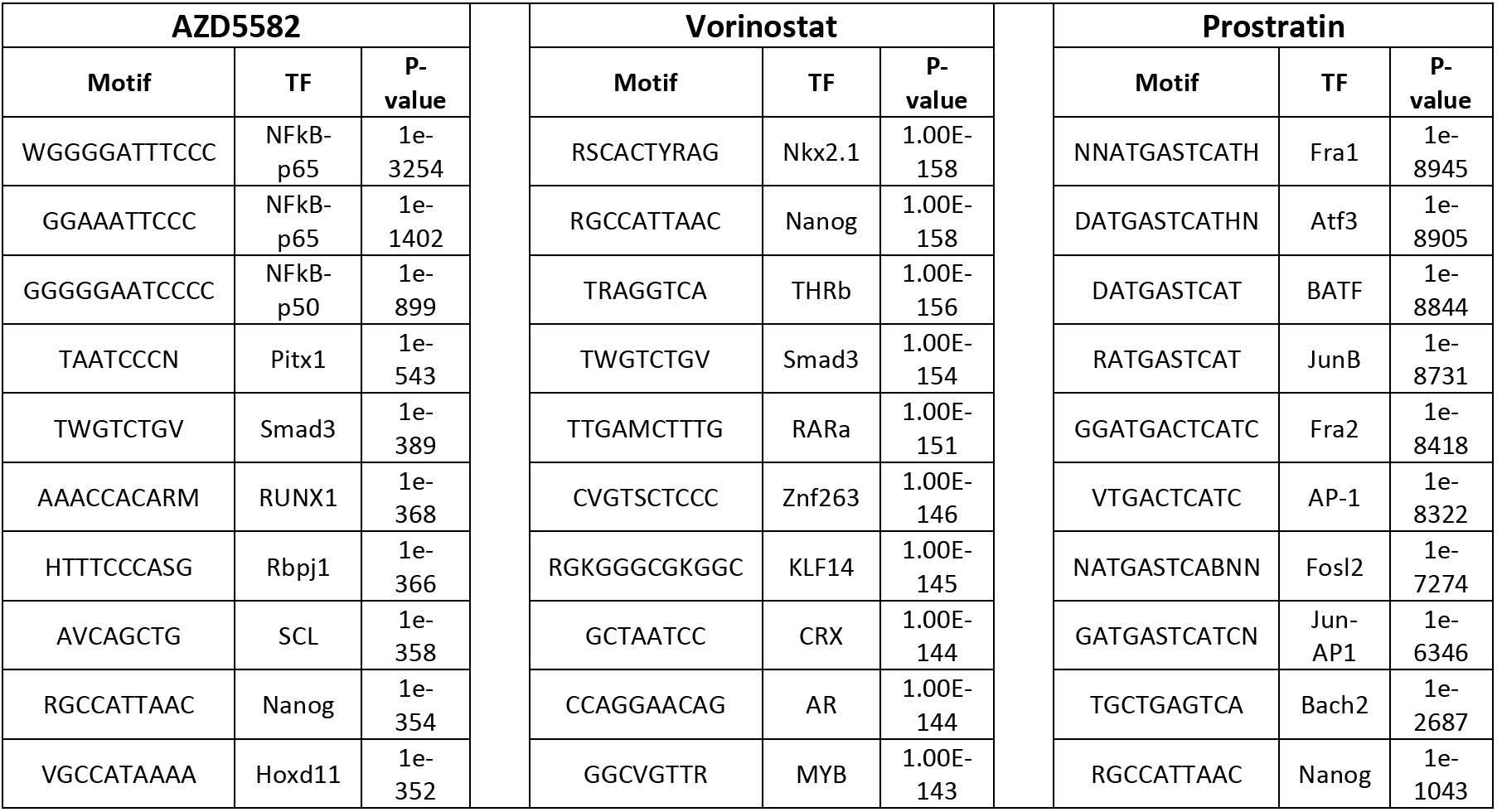
TF motifs with increased enrichment after LRA stimulation. The top 10 most significantly increased TF motifs in accessible chromatin after stimulation of infected with LRAs are shown. Also shown are putative TF binding partners for those motifs and the P-value of significance for enrichment. Data are derived from four independent experiments.

### LRAs induce opening and closing of TF binding sites

To examine whether LRA stimulation triggers changes in the accessibility of bindings sites for specific TFs, we also performed HOMER analysis on the differentially open peaks after LRA stimulation. Each LRA induced a distinct set of TF binding sites to open after stimulation (**Tables S3–5**). For AZD5582, the most strongly enriched TF binding site was NF***κ***B, consistent with this compound’s known ability to trigger non canonical NF***κ***B activation (Nixon et al., 2020). Other AZD5582-induced TF binding sites included PITX1, SMAD3 and RUNX1. For vorinostat, the most highly induced TF binding site was NKX2.1, followed Nanog, THRb and SMAD3. Prostratin induced the opening of binding sites for numerous TFs, most potently for members of the AP-1 family (BATF, FRA1, FRA2, JUNB), suggesting that activation of AP-1 could be a key mechanisms of latency reversal for prostratin and other PKC agonists.

### CTCF regulates HIV latency

We also examined enrichment of TF binding sites present in the set of differentially open peaks of GFP+ and GFP-cells through a different approach – using the overall genome as a background reference baseline (**Table 3**). Although many of the TFs binding site motifs identified by this approach were also identified by our previous analysis, several notable differences were apparent. Interestingly, the transcription factor with the highest degree of enrichment in the differentially open peaks between GFP+ and GFP-cells relative to the overall genome was the insulator protein CTCF. An overall set of 366 CTCF binding sites were differentially accessible between these populations, with 306 CTCF binding sites more accessible in GFP+ cells, and 60 sites more accessible in GFP-cells (**Figure 6A**). Furthermore, CTCF binding sites were also highly represented within differentially accessible peaks after LRA stimulation (**Figure 6B**). These data suggest that widespread changes in accessibility for CTCF binding sites are associated with HIV latency and latency reversal. These changes occur in both directions, although more CTCF sites were accessible during active infection than in latency. CTCF is a large, ubiquitously expressed regulator of gene expression with an 11 zinc finger central domain and N and C termini that interact with numerous protein partners (Ghirlando and Felsenfeld, 2016). The human genome contains 15,000-40,000 CTCF binding sites (Ghirlando and Felsenfeld, 2016; Holwerda and de Laat, 2013), and CTCF homodimerization drives the formation of large topologically-associated DNA domains (TADs) by mediating chromatin loop structures (Chang et al., 2020). These loops serve to insulate domains of transcriptional regulation by disconnecting genes from nearby enhancer elements, and by restricting lateral spread of domains of both inhibitory and activating histone modifications (Hansen et al., 2019; Saldaña-Meyer et al., 2019). Interestingly, CTCF is known to play an important role in regulating transcriptional latency for a number of other viruses, including HSV, KSHV and HPV (Chau et al., 2006; Washington et al., 2018). However, CTCF is not known to have a role in regulation of HIV latency. To investigate the possible role of CTCF in HIV latency we examined the impact of CTCF depletion on viral gene expression in HIV latency models. First, we transduced N6 cells, a latently infected T cell line model that contains a transcriptionally silent HIV reporter virus that encodes murine heat shock antigen (HSA) (Bradley et al., 2018), with lentiviruses that express an shRNA targeting CTCF or a scrambled control shRNA. Western blot analysis of CTCF confirmed specific reduction in CTCF expression in the cells transduced with the CTCF-targeting shRNA (**Figure 7A**). At two weeks post transduction, we measured viral reactivation by HSA expression on the surface of the transduced cells using flow cytometry. Notably, we observed potent reexpression of HSA in the CTCF-depleted cells, indicating that CTCF expression is required to maintain repression of HIV transcription in this cell line (**Figure 7B, C**). We next examined the impact of CTCF depletion in our primary CD4 T cell latency model. We activated CD4 T cells through TCR stimulation for 48h, then co-infected with HIV-GFP. At 3dpi, we purified infected (GFP+) cells, then nucleofected the infected cells with Cas9/sgRNA ribonucleoprotein complexes targeting the viral protein Tat, CTCF or with a non-targeting control (**Figure 7D**). Quantitative PCR one week after nucleofection confirmed roughly 50% reduction of CTCF expression in the targeted cells (**Figure 7D**). The infected cells were then cultured for four weeks and the percentage of latently infected (GFP-) cells over time measured by flow cytometry (**Figure 7F**). Tat targeted cells, as expected, demonstrated a more rapid decline in viral gene expression, confirming the sensitivity of this system to perturbations in regulators of viral transcription (**Figure 7F**). Notably, CTCF depleted cells exhibited a reduced level of HIV latency beginning at 2wpi, and this effect became more prominent at 3wpi and 4wpi. Overall these results demonstrate a role for CTCF in the establishment or maintenance of HIV latency, and highlight a previously unknown mechanism of transcriptional repression for HIV.

**Table 3:**
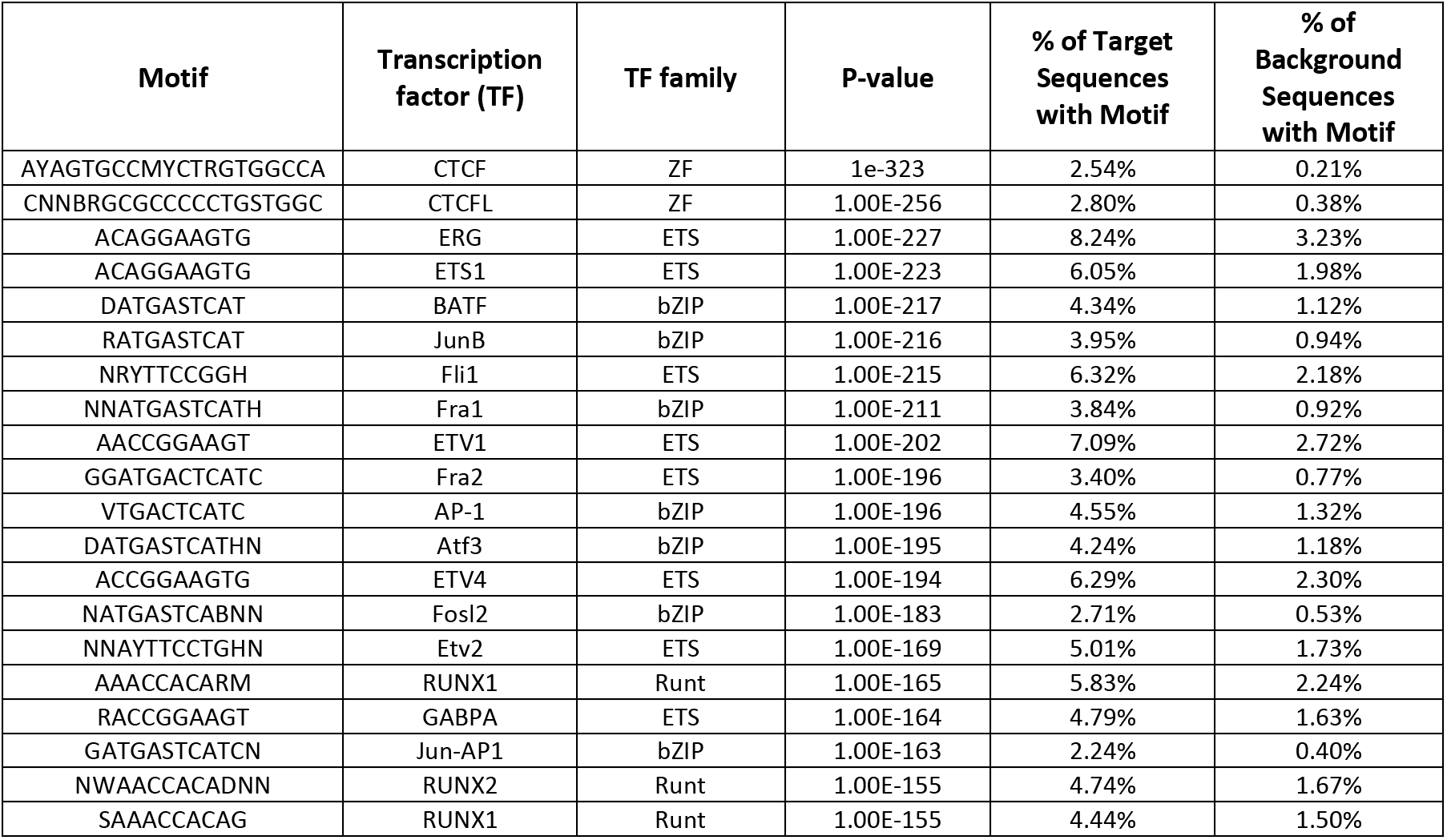
TF motifs enriched in differentially open peaks compared to overall genome. HOMER motif enrichment analysis of all differentially open/closed peaks between GFP+ and GFP-infected cells, using the whole human genome as a reference baseline. The 20 most enriched sequences and the putative TFs that bind these sequences are displayed.

**Figure 6:**
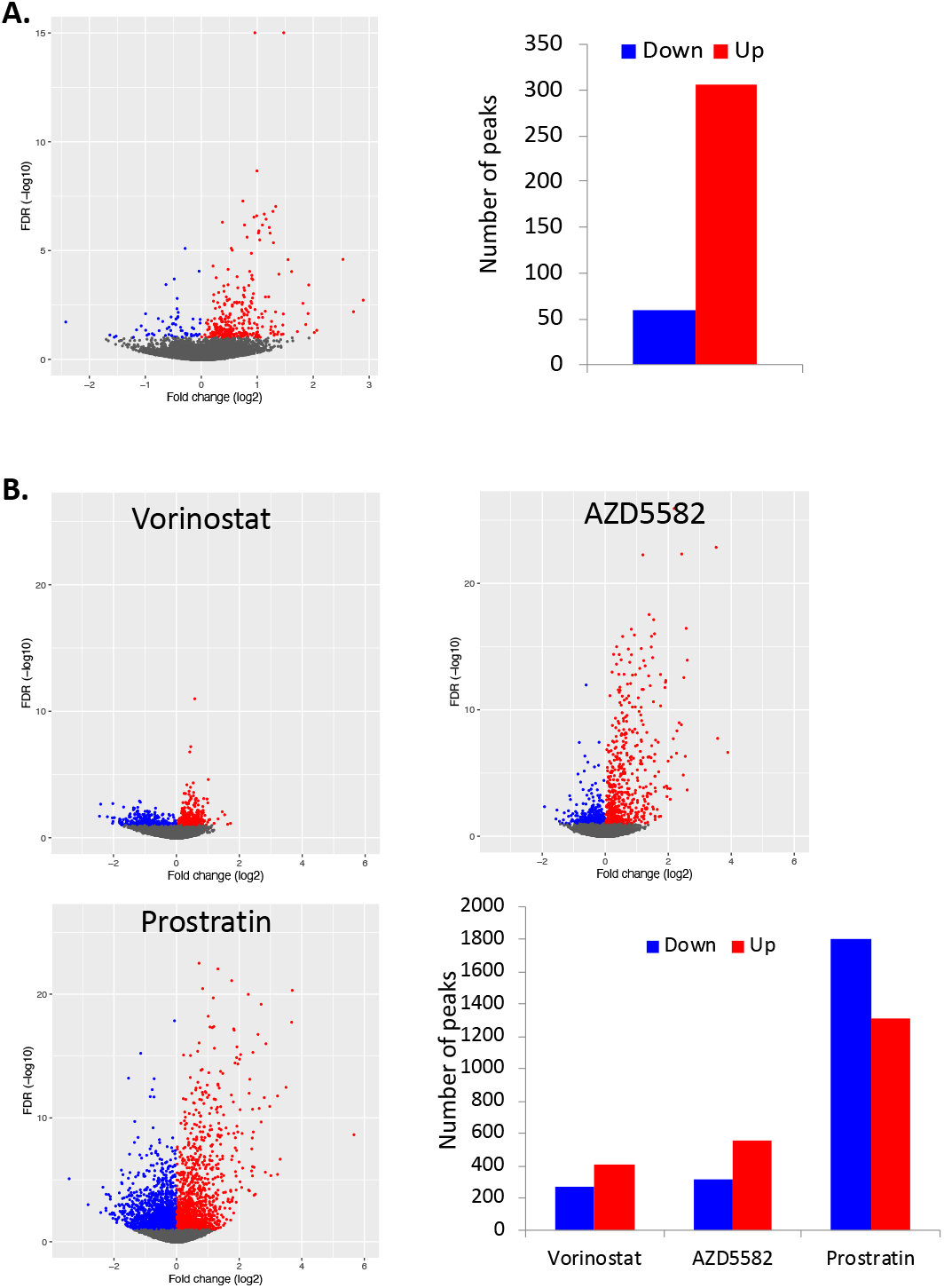
Differential opening of CTCF sites in latency vs active infection, and during latency reversal. **A.** Differentially accessible regions of the cellular genome containing consensus CTCF binding sites were identified by comparison of actively infected (GFP+) and latently infected (GFP-) cells, and fold changes versus FDR values are displayed as a volcano plot. Red datapoints indicate regions that are more accessible in actively infected (GFP+) cells (“up”), while blue datapoints represent regions that are more open in latently infected (GFP-) cells (“down”). **B**. Host cell chromatin peaks containing CTCF binding sites are displayed for the three LRA drug treatment conditions. Peaks with significantly different accessibility (FDR <0.1) after stimulation of HIV infected cells are highlighted in red (upregulated by LRA) and blue (downregulated by LRA). The data represents an aggregate of infected cells from four independent replicate experiments.

**Figure 7:**
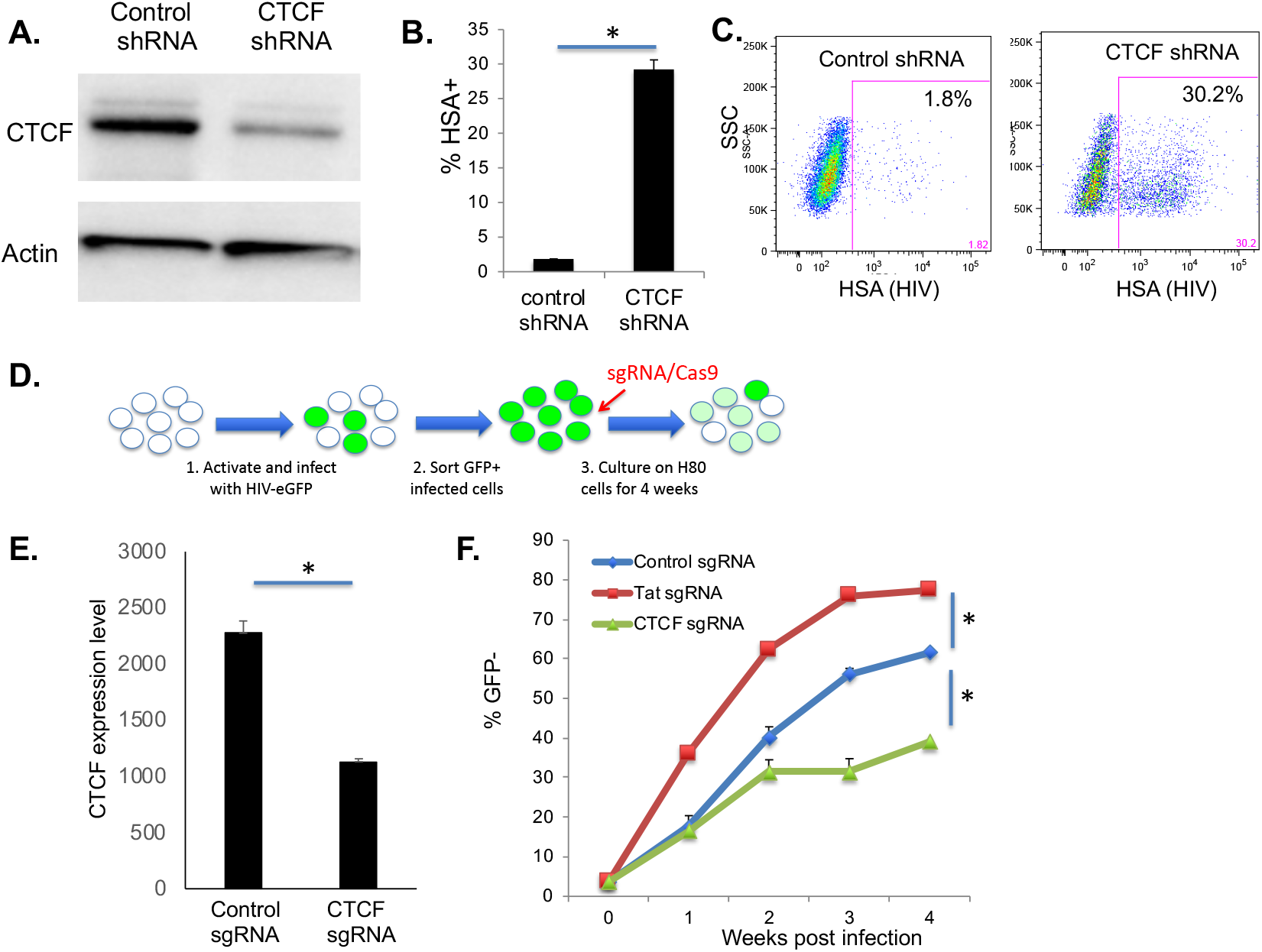
CTCF depletion in latently infected T inhibits HIV latency. 2D10 cells or N6 cells were transduced with lentiviruses that express mCherry and an shRNA targeting CTCF or a control scrambled shRNA. **A.** Transduced N6 cells were analyzed western blot to examine CTCF expression as well as a loading control (β-Actin). **B.** Transduced N6 cells were analyzed at 2 weeks post infection by flow cytometry, and the percentage GFP+ cells within the mCherry+ gate determined. Data represent the average of two technical replicates from one of two representative experiments. **C**. A representative FACS plot of HSA expression in shRNA transduced N6 cells. **D**. Primary CD4 T cells at 2 days post activation, were infected with HIV-GFP, then nucleofected with Cas9 complexed with a control sgRNA, a Tat sgRNA or a CTCF sgRNA. **E.** At 1 week post infection, the impact on CTCF mRNA levels, relative to beta-actin was quantified by Taqman qPCR. CTCF expression is displayed in arbitrary units and represents the average of four replicates. **F**. The infected/nucleofected cells were analyzed for GFP expression over four weeks of infection and the percent of latently infected (GFP-) cells was determined. Each datapoint is the average of two independent nucleofection reactions. Asterisks represent significant differences (P<0.05).

## Discussion

The presence of a latently infected reservoir is recognized as a primary barrier to curing HIV infection (Margolis, 2014). As such, the mechanisms by which latent infection is established and maintained are of critical importance. By characterizing these mechanisms we may identify strategies to either induce viral gene expression leading to infected cell clearance (“shock and kill)”, or induce permanent silencing of the proviruses (“block and lock”) (Mousseau and Valente, 2015; Siliciano and Greene, 2011). Viral gene expression is regulated by host cell transcription factors and epigenetic mechanisms. Dynamic changes and cell-to-cell variation in these factors likely leads to specific cellular environments that favor or disfavor latency. However, viral transcription can also be influenced by stochastic fluctuations in the Tat transcription factors early in infection (Razooky et al., 2015; Weinberger et al., 2005). Nevertheless, the ability of LRAs targeting host cell factors to induce viral reactivation in both animal models and in human clinical studies demonstrates the value of targeting host cell pathways to manipulate latency (Archin et al., 2012; Nixon et al., 2020; Søgaard et al., 2015). Thus, latency initiation and reversal both likely result from a combination of stochastic and deterministic processes intrinsic to both the virus and the host cell. These processes yield a diverse population of persistently infected cells, with intermittent viral expression and re-entry into quiescence, undergoing both proliferation and death,

The rare nature of latently infected cells and the lack of markers for their purification, make direct observation of cellular events in infected cells from patients extremely difficult. In this regard, model systems have been invaluable in the discovery and validation of new targets. In this study, we have discovered, using a primary cell model system, that viral latency is a stable, heritable phenotype that is rapidly re-established after stimulation and transmitted to daughter cells during cell division. These findings expand the understanding of epigenetic mechanisms likely contribute to the maintenance of latency through cell division, an observation that has significant implications for the maintenance of the HIV reservoir *in vivo* through clonal expansion. Epigenetic memory of HIV latency through cell division and T cell development are likely to explain the presence of expanded latently infected clones within diverse T cell subset identities present in infected individuals on therapy. The rapid and repetitive re-establishment of latency after stimulation also has implications for clearance strategies against reactivated latently infected cells *in vivo*. These results suggest that a temporally limited window of antigen presentation will likely occur after LRA dosing before latency is reestablished and antigen expression wanes.

By analysis of infected cells with ATACseq, we have also discovered that HIV latency is preferentially established in cells that adopt a specific global chromatin structure after activation and return to a resting state. This finding suggests that latency is intricately connected to cell-intrinsic chromatin dynamics, and that these dynamics could represent an important target for preventing latency establishment, reversing latency once established, and preventing a return to latency following latency reversal. The specific mechanisms by which different patterns of overall chromatin accessibility are regulated in these infected cells as latency is established are unclear, and will require further investigation. We have also demonstrated that accessibility of the viral genome is significantly reduced in latently infected cells relative to actively infected cells. This finding confirms previous results in cell line models showing the appearance of restrictive chromatin structures associated with entry of HIV into latency (Pearson et al., 2008). The closed nature of the provirus likely impedes recruitment of transcription factors essential for initiating viral transcription, and processive RNA polymerase transcription, thereby contributing to latency. Notably, three LRAs with distinct mechanisms of action caused significant re-opening of the provirus, indicating that re-opening of HIV is likely a key feature of latency reversal by LRAs.

This study also provides new insight into the mechanisms of latency by identifying a set of host cell TFs whose activity correlates with viral gene expression. Specifically, by comparing the differentially open regions of chromatin in the latent or actively infected cells to the overall set of open regions across infected CD4 T cells, active viral expression was correlated with elevated activity of AP-1, GATA, and RUNX TFs, while latency was associated with elevated activity of Forkhead (FOX) and Kruppel-like factor (KLF) TFs (**Figure 8**). Importantly, this list contains several TFs that have well-established roles in regulating HIV transcription, indicating the biological relevance of the findings. AP-1, for example has long been identified as a regulator of HIV gene expression (Duverger et al., 2013; Roebuck et al., 1996; Yang et al., 1999). AP-1 is a family of heterodimeric basic leucine zipper TFs that recognize a consensus binding site (TGA G/C TCA), and are potently activated by stimuli that promote HIV gene expression, such as phorbol esters. There are several AP-1 binding sites in the HIV LTR, and these have been shown to be important for HIV transcription and replication, suggesting that the correlation between AP-1 activity and viral gene expression in our model is likely through direct activity of AP-1 at the HIV LTR. Interestingly one AP-1 site has recently been shown to regulate latent infection (Duverger et al., 2013). GATA TFs have also previously been identified as regulators of HIV transcription. GATA3, in particular, has been shown to bind the HIV LTR and promote HIV gene expression (Yang and Engel, 1993). Notably, we have previously reported that elevated GATA3 transcript levels correlates with active HIV transcription in this model system (Bradley et al., 2018). From the list of TFs with enriched activity in latently infected (GFP-) cells, FOXP1 was also previously reported by our group as exhibiting elevated expression in latently infected CD4 T cells (Bradley et al., 2018). This TF is associated with a quiescent central memory T cell (Tcm) phenotype, consistent with the hypothesis that cells that adopt a quiescent Tcm phenotype are more prone to latent infection (Garaud et al., 2017). Interestingly, a related Forkhead TF, FOXO1, has also recently been reported to promote HIV latency (Roux et al., 2019; Vallejo-Gracia et al., 2020). In addition to these known HIV-regulating TFs, several TFs with no known role in HIV transcription were identified by our study, and defining their role in latency will likely lead to the discovery of new mechanisms of transcriptional repression for HIV.

**Figure 8:**
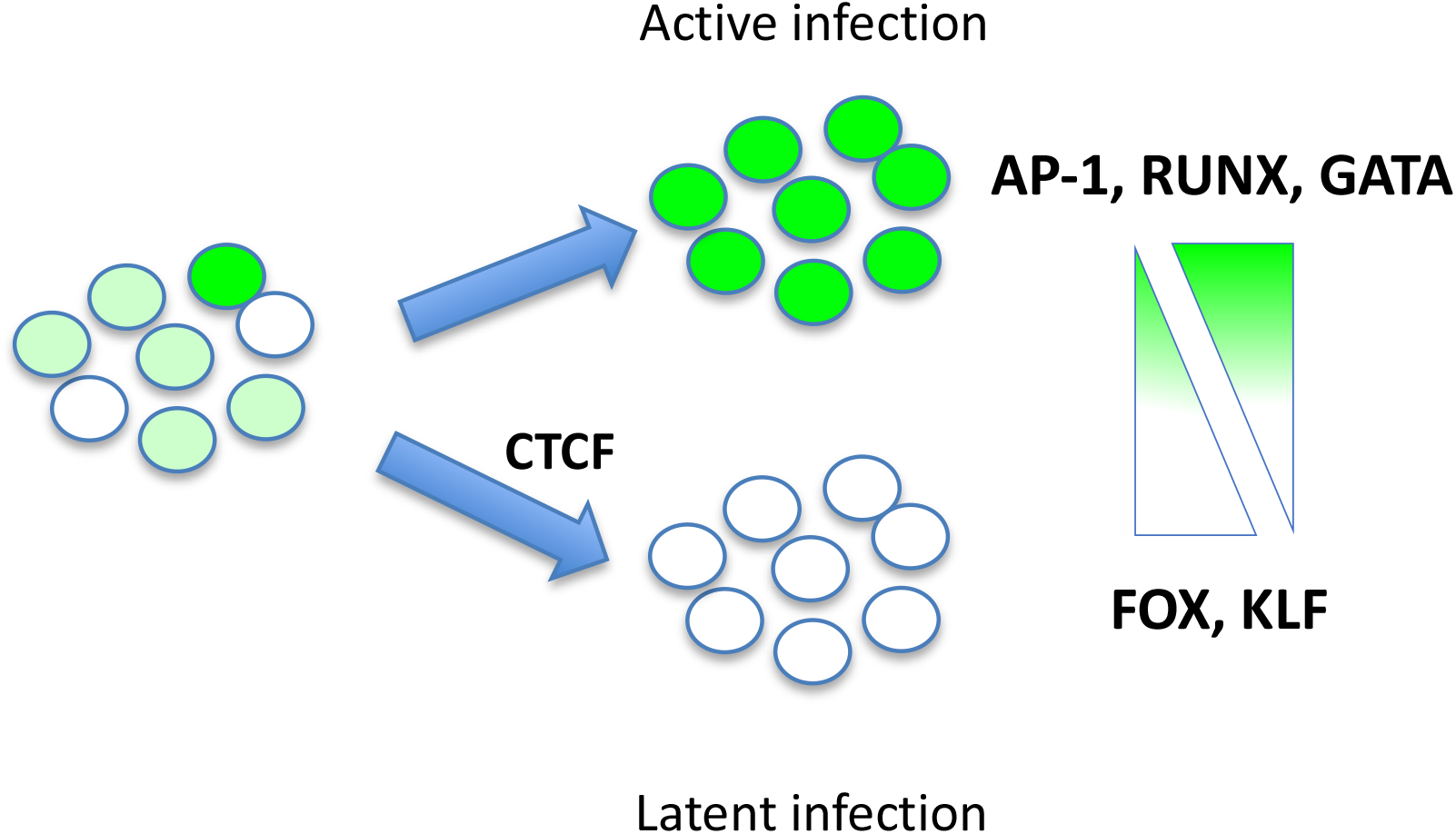
Schematic model of transcription factor activity and HIV latency. HIV-GFP infected cells exhibit variegated levels of transcriptional silencing. Comparison of actively infected (green) and latently infected (white) CD4 T cells reveals latency associated TF activity. Latently infected cells have elevated Forkhead (FOX) and Kruppel-like factor (KLF) TF activity, and reduced AP-1, RUNX and GATA TF activity relative to actively infected cells. The chromatin insulator CTCF promotes HIV latency.

We also examined enrichment of TF motifs in the differentially open chromatin of infected cells relative to the overall human genome. Interestingly, by this comparison, the differentially open chromatin regions of infected CD4 T cells were highly enriched with binding sites for CTCF, indicating that possible dynamic changes to chromatin looping may distinguish latently infected cells from actively infected cells. Furthermore, by selective knockdown of CTCF in a T cell line model of latency (N6) and in a primary cell latency model, we demonstrated that CTCF expression is required for maintaining HIV silencing in HIV infected T cells. These results thus identify CTCF as a novel regulator of HIV latency. Notably, previous studies have shown that CTCF plays a key role in regulating latency for herpesviruses (Chau et al., 2006; Washington et al., 2018). In these cases dynamic CTCF binding to cognate sites within the viral genomes directly regulates switching between lytic and latent gene expression programs. It is unknown if CTCF binds directly to the HIV genome. As such, the precise mechanism by which CTCF regulates HIV latency is unknown and will require further investigation. CTCF has several known molecular functions that could potentially affect HIV gene expression. In particular, CTCF binding to DNA mediates chromatin looping and the formation of topologically associated domains (TADs), and serves to insulate genes from adjacent enhancer elements and zones of histone modifications (Chang et al., 2020). It is conceivable that, in many infected cells, HIV gene expression is impacted by insulation of the HIV promoter from nearby enhancer elements and that disruption of the insulator function of CTCF allows longer-range promoter/enhancer interactions to occur that promote viral reactivation. Alternatively, many latent proviruses may be trapped in TADs that are enriched for repressive histone modifications such as H3K9me3 and H3K27me3, and removal of the CTCF boundary allows replacement of these heterochromatic histone modifications with activating marks. CTCF can also regulate sub-nuclear localization of chromatin regions and mediate interactions with the nuclear lamina (Fiorito et al., 2016). As such, it is also possible that CTCFs ability to regulate latency is determined by promoting spatially restricting access of the provirus to activating transcription factors within the nucleus.

It is also unclear what might regulate differential binding activity of CTCF within our latency model system. CTCF is ubiquitously expressed in mammalian cells, and CTCF transcripts were not differentially expressed between the latently infected (GFP-) and actively infected (GFP+) cells. It is possible that CTCF binding is regulated by expression or differential binding of an interacting partner. CTCF contains three major structural domains – a central DNA binding domain with 11 zinc fingers, and long N and C terminal domains. All three domains interact with an array of binding partners, including Y-box binding protein 1 (YBX1), Chd4 and Snf2h (Ghirlando and Felsenfeld, 2016). CTCF itself can be regulated by phosphorylation, sumoylation and ADP ribosylation (Del Rosario et al., 2019). CpG methylation of CTCF target sites can also regulate CTCF binding (Wang et al., 2012). It is also possible, given CTCFs fundamental role in transcriptional regulation, that CTCFs role in repressing HIV transcription is indirect, and mediated through an as yet unidentified factor whose expression or activity is regulated by CTCF. Deciphering the precise mechanism behind CTCFs role in latency will need to be clearly defined. No therapeutic strategies for targeting CTCF currently exist, but the results we describe in this manuscript suggest that inhibition of CTCF activity or a pathway regulated by CTCF could serve as a strategy for impacting HIV latency clinically.

## Methods

### Viruses and Cell culture

Stocks of HIV-GFP were generated by co-transfection of 293T cells with the pNL4-3-Δ6-dreGFP plasmid (generous gift of Robert Siliciano) and the packaging plasmids PAX2-GagPol) and MD2-VSVG, using Mirus LT1 reagent. At 2 days post transfection virus was harvested from the supernatant and clarified by low speed centrifugation, followed by filtration through a 45uM filter. shRNA expressing lentivirus plasmids targeting CTCF were purchased from Genecopeia (HSH060478-LVRU6MP) and viral particles generated as for HIV-GFP. Latently infected N6 cells are a clone of Jurkat cells that contain a full length clone of the NL4-3 strain of HIV in which the Nef open reading frame has been replaced by an HSA-P2A-Luciferase cassette (generous gift from David Irlbeck, Viiv Healthcare).

### Primary cell latency model

Primary CD4 T cells were isolated from fresh whole blood (purchased from Gulf Coast Regional Blood Center) by Ficoll isolation of peripheral blood mononuclear cells (PBMCs), then magnetic negative selection of CD4 T cells using a CD4 enrichment kit (Stem Cell). Total primary CD4 T cells were activated using anti-CD3/CD28 beads (Thermo Fisher) at a ratio of one bead per cell for 2 days, then infected with HIV-GFP viral supernatant by spinocculation for 2hrs at 600g with 4ug/mL polybrene. The infected cells were then resuspended in fresh RPMI with IL-2 (Peprotech) at 100U/mL and incubated for two days before actively infected (GFP+) cells were sorted using a FACSAria flow sorter (Becton Dickson). The purified infected cells (GFP+) were then maintained at 1-2M/mL with fresh media and IL-2 added every 2-3 days for 2 weeks. From 2wpi, cells were co-cultured with H80 cells (provided by D Bigner, Duke University) in the presence of 20U/mL IL-2 for up to 12wpi. During co-culture the cells were fed with fresh media and IL-2 every 2-3 days, and moved to a fresh flask of H80 feeder cells once a week. For latency reversing agent stimulation, AZD5582, vorinostat and prostratin were generous gifts from David Irlbeck (Viiv Healthcare), and were reconstituted in DMSO at 10mM, before dilution into media to working concentrations of 250nM.

### ATACseq library construction and sequencing

Infected cultures of CD4 T cells were stained with the viability dye Zombie Violet (ZV, Biolegend) and 50,000 live (ZV-) cells were sorted into 5mL polypropylene tubes using a FACSAria flowsorter (Becton Dickson) per replicate. The sorted cells were then washed with phosphate buffered saline (PBS) and resuspended in 50ul TD buffer (Illumina) with 0.01% digitonin and 2.5ul TDE Tagmentation enzyme (Illumina). The cells were then agitated in a thermomixer at 37C, 300rpm for 30mins to allow transposon-mediated tagmentation to occur. Tagmented DNA was then purified a Minelute purification kit (Qiagen). DNA fragments were PCR amplified using dual barcoded primers and repurified using a Minelute kit followed by a 1.5x final cleanup step with Ampure SPRI beads (Agencourt). Libraries were quantified by Qubit DNA assay (Invitrogen) and size distribution of the library was determined using a High Sensitivity DNA BioAnalyzer chip (Agilent). Samples were sequenced using an Illimuna Hiseq2500 to a depth of 50-150M reads with 150bp paired end sequencing.

### Bioinformatic Analysis pipeline

Sample fastq files were combined by read end and validated with FastQC v0.11.8. Combined fastq read end pair files were aligned with BWA_MEM v0.7.17 (Li, 2013) against the GRCh38 genome with an added HIV chromosome sequence. The BWA_MEM -Y flag was used to support soft clipping and alignment across HIV (non-clonal) integration sites. Aligned bam files were sorted and optical duplicates were flagged with Picard Tools v2.21.1. Picard tools and SAMtools v1.9 (Li et al., 2009) were used to verify the quality of the alignments. Filtering prior to read counting was done by SAMtools to remove duplicate, unmapped, and unpaired reads. HIV-aligned reads were duplicated into separate bam files and merged by group with SAMtools for visualizing in IGV (Thorvaldsdóttir et al., 2013). Read counting was done using csaw v1.16.1 (Lun and Smyth, 2016), with parameters set to count reads with map quality greater than or equal to 10 and inferred fragment length less than or equal to 2000. Read ends were counted separately and combined into bins of 73 bp windows (~1/2 nucleosome width) tiled across the entire genome. Reads were also counted into 5000 bp bins, generating a second count matrix at lower resolution. The csaw function filterWindows() was used to calculate the log2 signal to noise of all 73 bp windows relative to the median of the large windows counts, scaled by relative window sizes. Low signal windows less than or equal to log2(2.5) over background were dropped. Per sample TMM normalization factors were calculated using the csaw normFactors() function, retaining potential global changes in accessibility. The normalized noise-filtered count matrices were analyzed by PCA using the prcomp() function of R 3.5. Review of the PCA results indicated the differential accessibility model for GFP+ vs GFP-data should include donor as a cofactor. For latency reversing agents treated cells, four replicate experiments were carried out using cells from a single donor and a simple treatment-control model for each. Differential accessibility was calculated using edgeR v3.23.4 (McCarthy et al., 2012), with global dispersion factors from estimateDisp(). Models were fit with glmQLFit() using robust= TRUE, and the significance calculated using glmQLFTest(). Finally, successive windows were merged into clusters across valleys of up to one bin width using mergeWindows() and combineTests() from csaw. The generated clusters from each modeled comparison were plotted as volcano plots visualizing the FDRs and the mean fold change (calculated as the mean fold change of all the cluster’s windows). Clusters were assigned category labels if they overlapped a matching HG38 annotated region, e.g. a “promoter” or “enhancer”, as provided by annotatr v1.8.0.

Transcription factor binding site enrichment for each modeled comparison was performed using findMotifsGenome.pl from HOMER v4.10 (Heinz et al., 2010). Input bed files were generated from the csaw cluster results, separately for the significantly up and significantly down clusters (FDR <= 0.1), and for the undifferentiated regions (FDR > 0.2). Using the included default set of transcription factors, HOMER was run to identify known TF binding sites enriched in the significant bed files relative to the undifferentiated background. Separate runs were performed examining the middle 50, the middle 200, and all bases of the significantly up and significantly down cluster regions. Results from the top hits were manually examined for relevant biological associations with HIV. Additionally, HOMER was used to compare the significantly different regions against the genome as a whole. A bed file was generated containing all clusters and analyzed with findMotifsGenome.pl using the -find parameter to locate all CTCF binding sites. Clusters overlapping CTCF sites were extracted and their differential accessibilities were plotted as volcano plots.

### Antibodies/Western blots

To extract protein, cells were washed in phosphate-buffered saline (PBS) then lysed in RIPA buffer (Sigma) supplemented with Complete protease inhibitors (Roche), before centrifugation to remove cell debris. Protein concentrations were then quantified using Bradford assay reagent (BioRad), and 40ug of protein per sample was run on a 4-12% Tris Glycine polyacrylamide gel (Invitrogen). The gel was then transferred to a PVDF membrane. The membranes were then incubated in Tris-buffered saline with 0.1% Tween 20 (TBST) with a primary detection antibody for 2-hrs, before washing with TBST. Primary antibodies used were anti-CTCF (clone D31H2, Cell Signaling) and beta-actin (ab49900, Abcam). Primary antibody staining was followed by incubation with an HRP-linked anti-rabbit antibody. Bands were the imaged by incubation of the membrane with ECL chemiluminescence reagents (ThermoFisher) and imaging on a ChemiDoc MP imager (BioRad). For flow cytometry for murine Heat Shock Antigen (HSA), cells were washed in PBS before staining with anti-HSA-APC (Biolegend) at 1:200 dilution for 30mins at 4C, before washing in PBS and analysis using a Fortessa flow cytometer (Becton Dickson).

### Quantitative PCR

Quantitative PCR for CTCF was performed using Taqman probes/primers (Thermo Scientific). Specific primer probe sets used were Hs00902016_m1 for CTCF, and Hs01060665_g1 for beta-actin. Reactions were run using primer/probes at 1:20 dilution and Taqman FastVirus one-step reaction mix (Thermo Scientific), and amplified using an Applied Biosystems QS3 cycler. Expression level was determined using cycle threshold relative to beta-actin and represented in arbitrary units.

### CRISPR/Cas9 nucleofection

CRISPR RNAs (crRNAs) targeting CTCF were pre-designed by IDT or designed with the BROAD institute GPP sgRNA designer. crRNAs and trans-activating RNAs (tracrRNA; IDT) were annealed in a thermocycler at a 1:1 ratio according to manufacturer’s instructions and stored at −20 C until use. Nucleofection experiments in primary CD4 T cells were performed six days post-infection using methods previously described for CRISPR nucleofection of either resting or activated human T cells (Oh et al., 2019; Seki and Rutz, 2018). CRISPR-Cas9 ribonucleoprotein (RNP) complexes were generated by mixing crRNA:tracrRNA complexes with ALT-R *S.p*. Cas9 nuclease V2 at a 3:1 molar ratio for 10 minutes at room-temperature. Infected CD4 T cells were washed with PBS and 3 million cells per condition were resuspended in 20uL buffer P2 (Lonza) with 4μM IDT electroporation enhancer. CRISPR-Cas9 RNP nucleofection was performed with the EH100 nucleofection protocol on a 4D Nucleofector device (Lonza) (Oh et al., 2019; Seki and Rutz, 2018). Immediately after nucleofection cells were resuspended in fresh prewarmed media containing 100U/mL recombinant human IL-2. sgRNA targeting sequences were: Non targeting: CGGAGGCTAAGCGTCGCAA, Tat: CCTTAGGCATCTCCTATGGC, CTCF: GAGCAAACTGCGTTATACAG

### Data accessibility

Underlying data files for ATACseq analysis available at https://webshare.bioinf.unc.edu/public/2020_HivLatency

## Acknowledgements

The authors gratefully acknowledge the following funding sources: NIH 1R01AI143381-01A1 (EPB), 1R61DA047023-01 (LIJ), and the UNC Center for AIDS research (CFAR). Research reported in this publication was supported, in part, by CARE, a Martin Delaney Collaboratory of the National Institute of Allergy and Infectious Diseases (NIAID), National Institute of Neurological Disorders and Stroke (NINDS), National Institute on Drug Abuse (NIDA) and the National Institute of Mental Health (NIMH) of the National Institutes of Health, grant number 1UM1AI126619. We also acknowledge assistance from the UNC flow cytometry core facility.

## Author contributions

Conceived the study: (EPB). Wrote the manuscript (EPB, SRJ, DMM) performed experiments (EPB, SB, SRJ, SS, JJP). Analyzed data (EPB, SRJ, AMT, LJM, JP).

## Supplemental data

**Fig S1:**
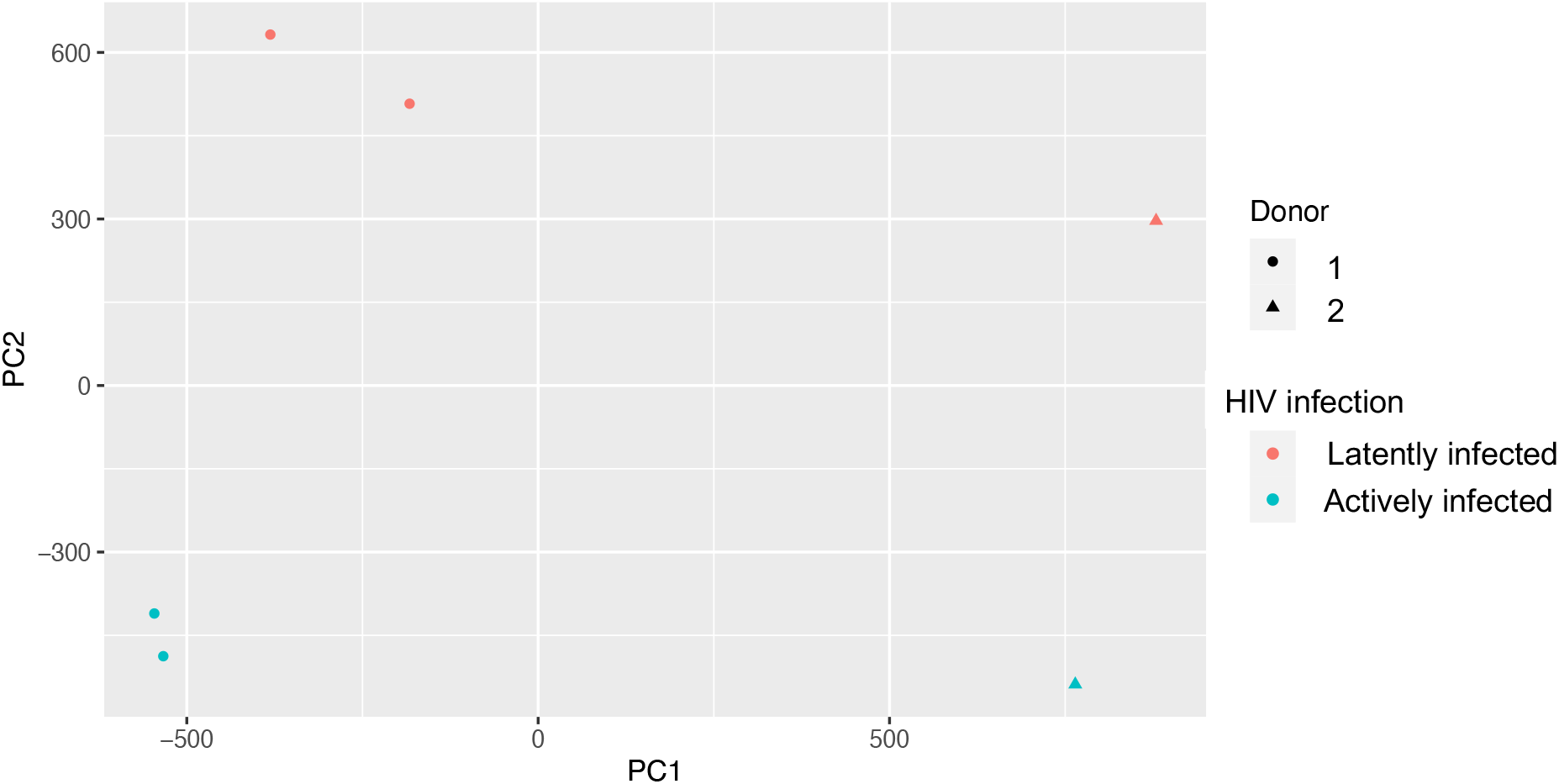
Principal component analysis of actively infected and latently infected cells ATACseq data. Principal component analysis was performed on ATACseq data from actively infected (GFP+, red datapoints) and latently infected (GFP-, blue datapoints) CD4 T cells. For Donor 1 (circles), two independent replicate experiments are shown. Donor 2 data represented by triangles.

**Table S1.**
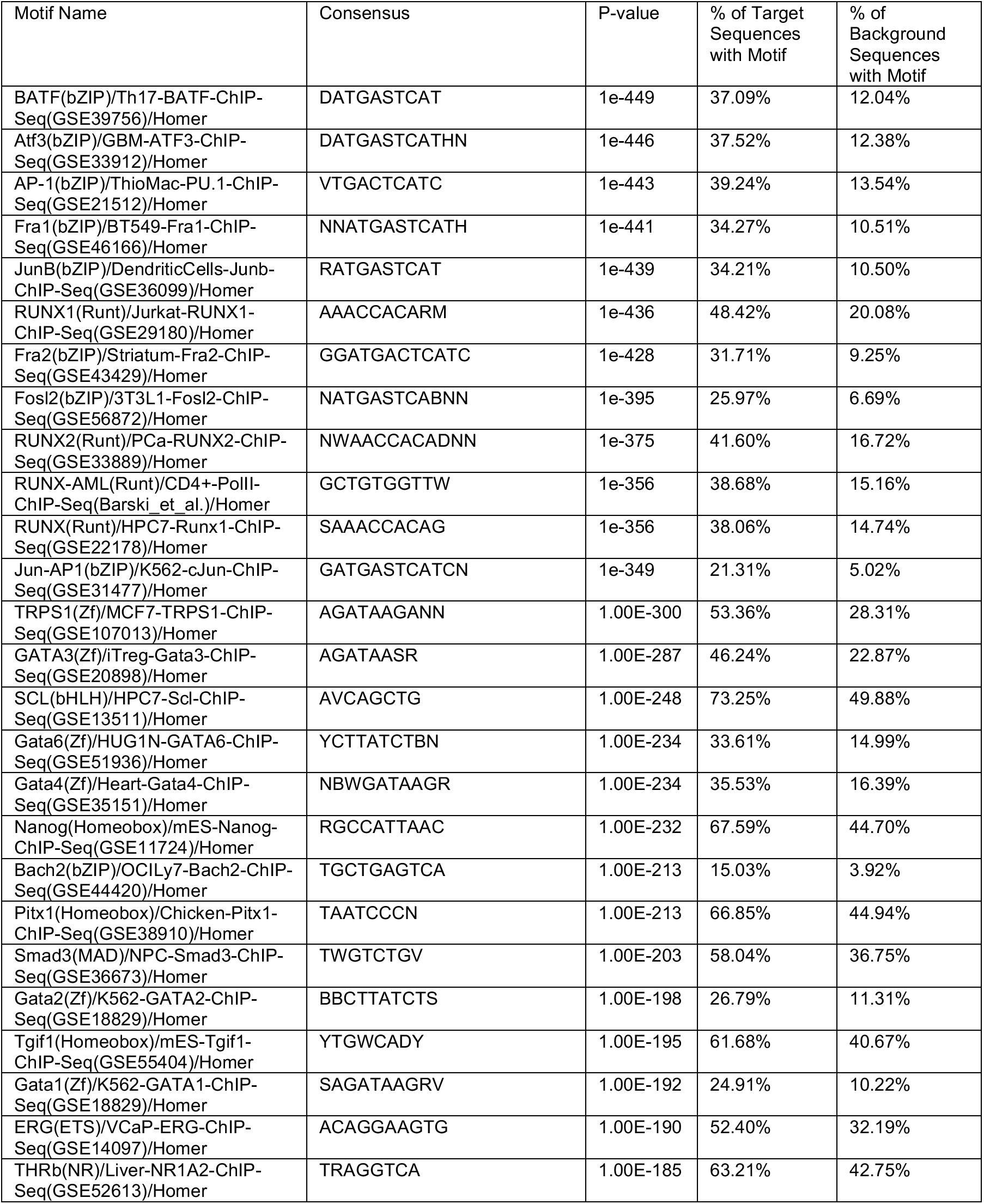
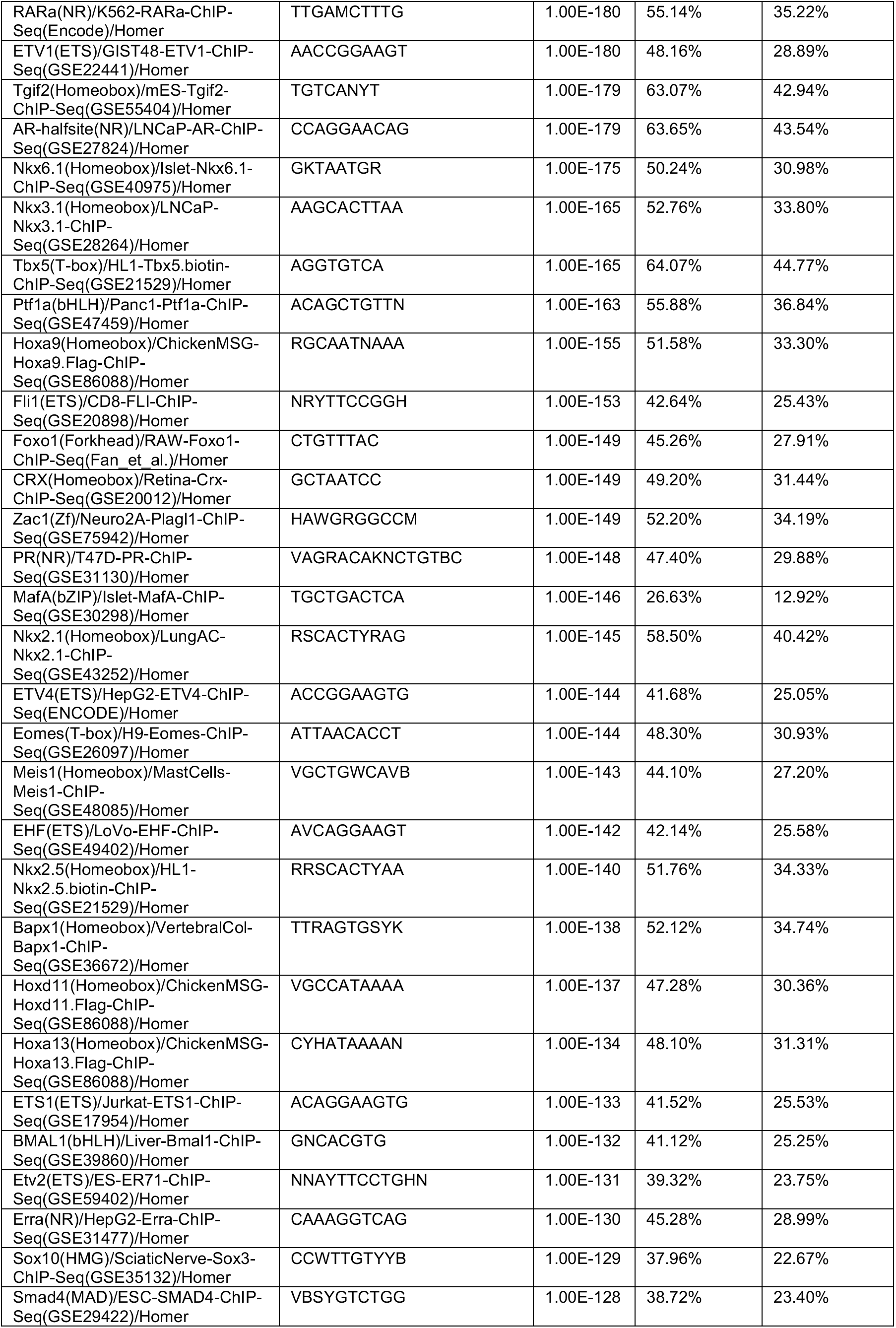
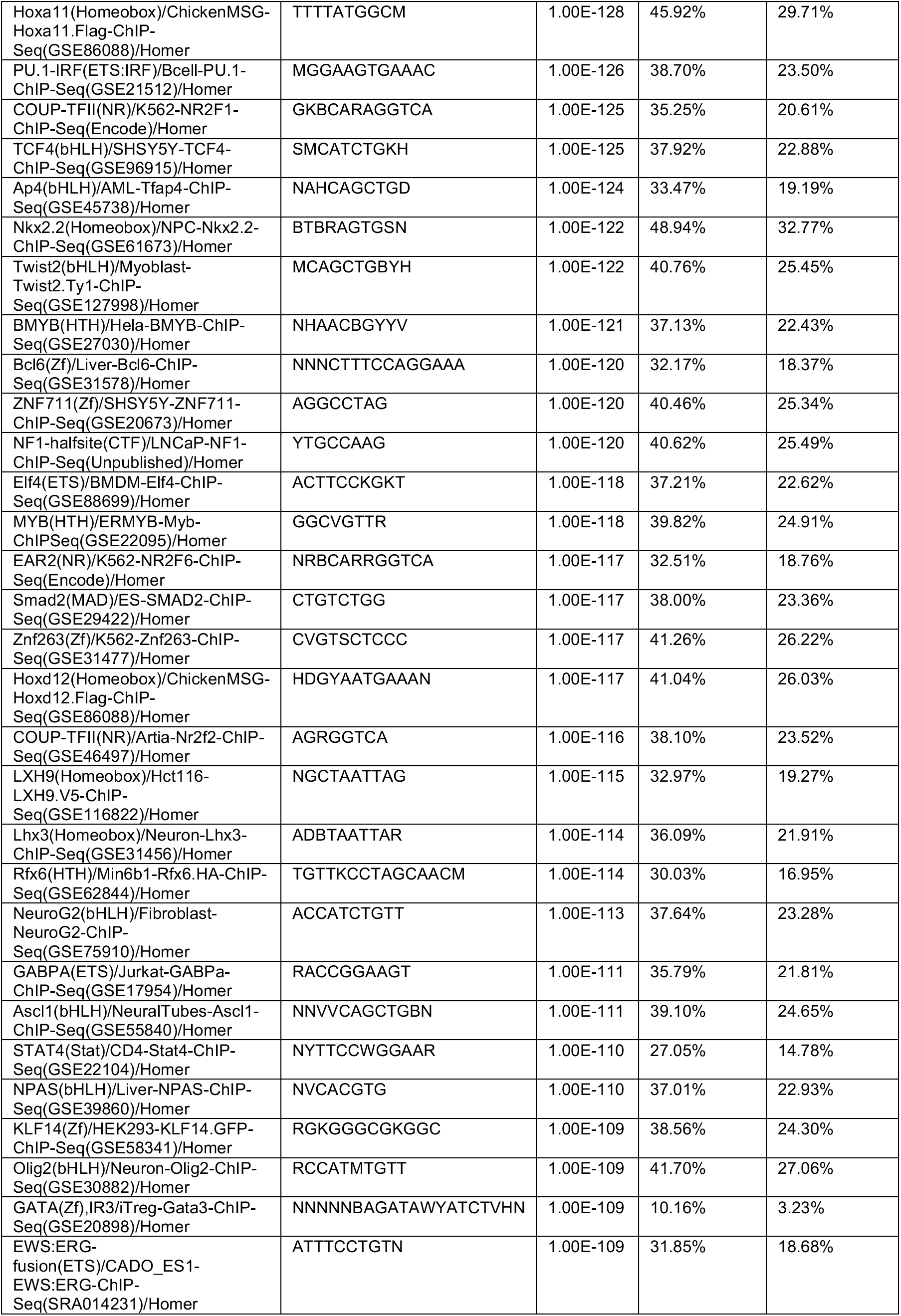
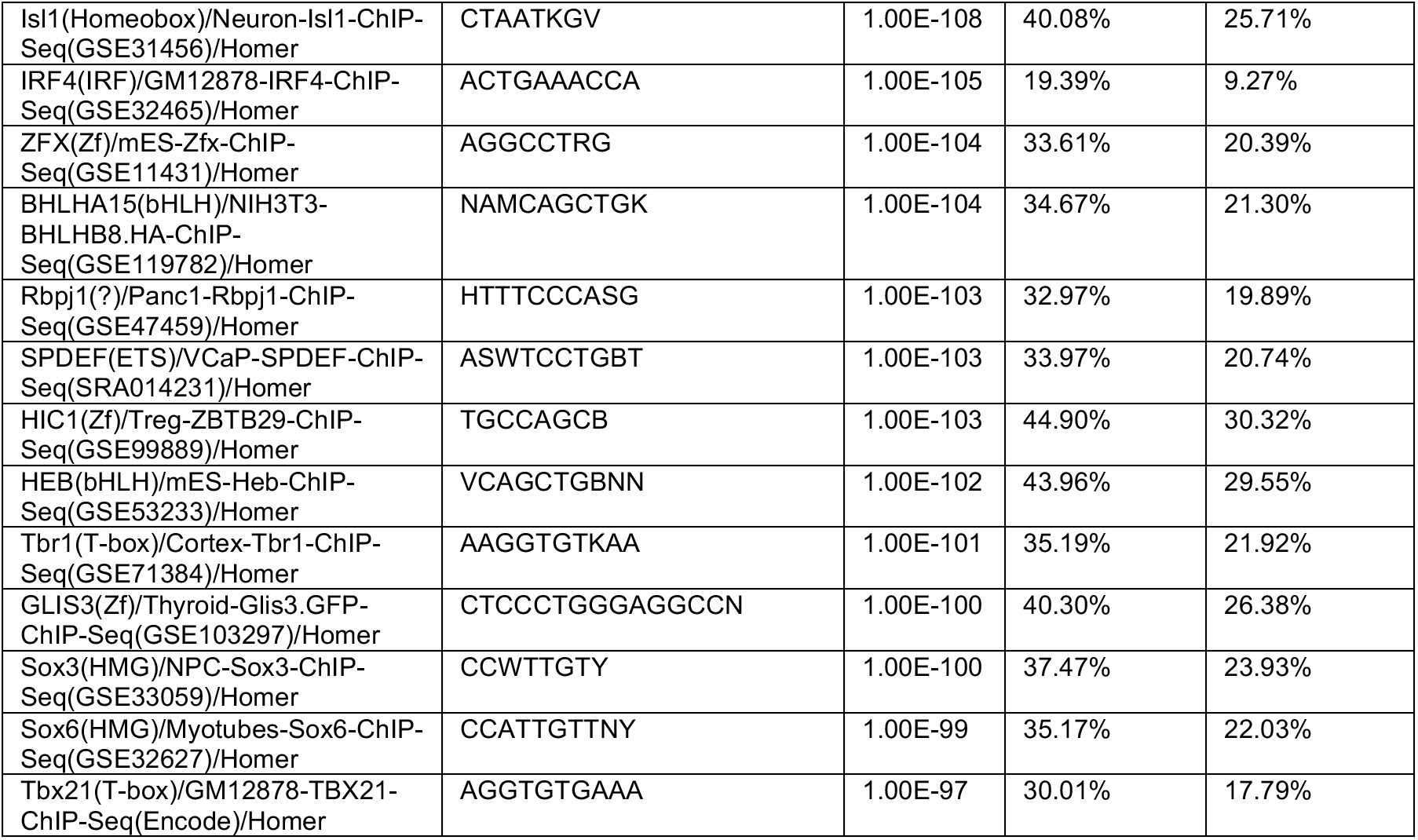
TF motif enrichment in actively infected cells. HOMER TF motif enrichment analysis of differentially open peaks between actively infected cells (GFP+) and latently infected (GFP-) cells. Top 100 enriched motifs in peaks preferentially open in actively infected cells are shown. Target Sequences represent genomic sequences that have significantly elevated accessibility in latently infected cells. Background Sequences represent all open chromatin regions in CD4 T cells.

**Table S2.**
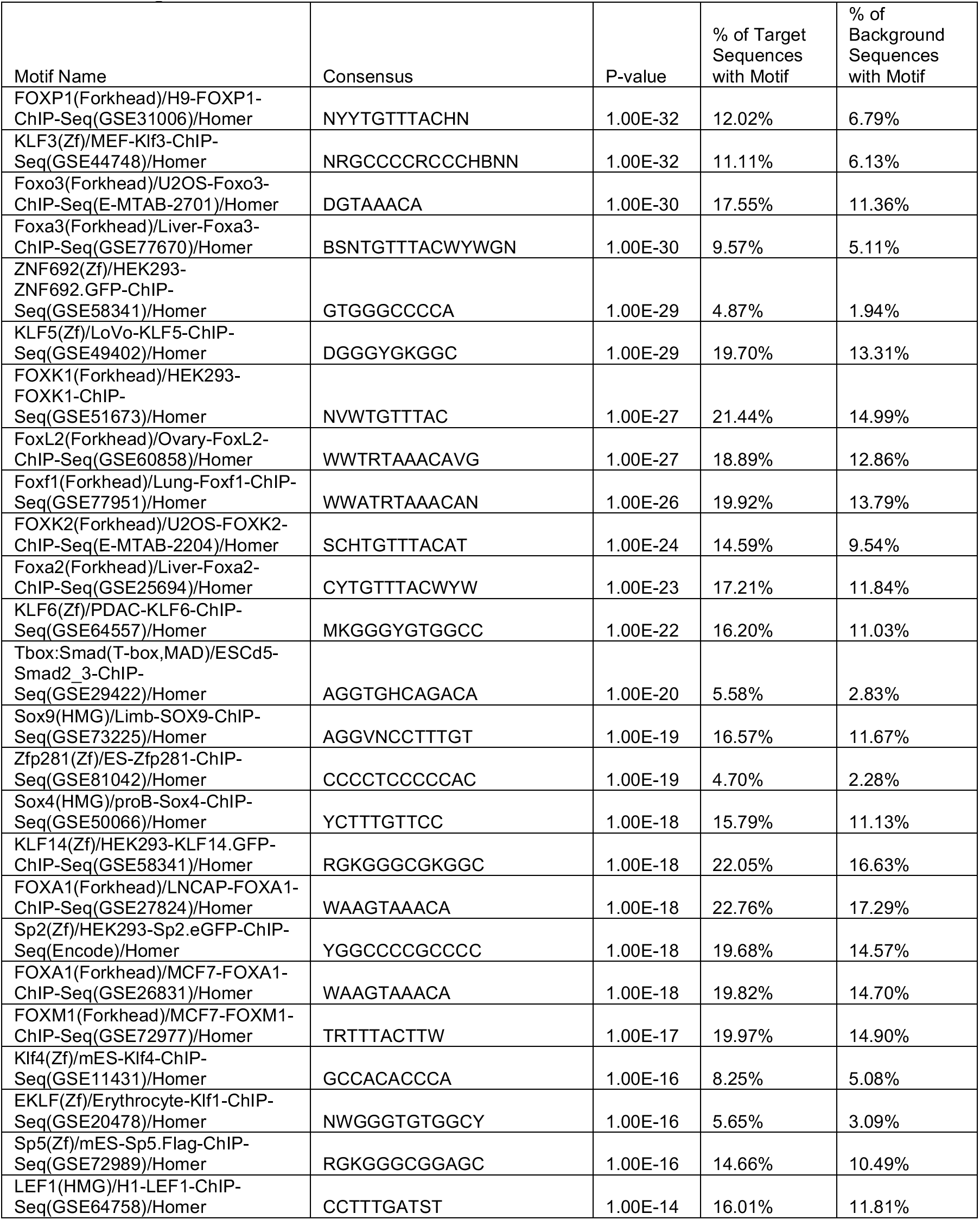
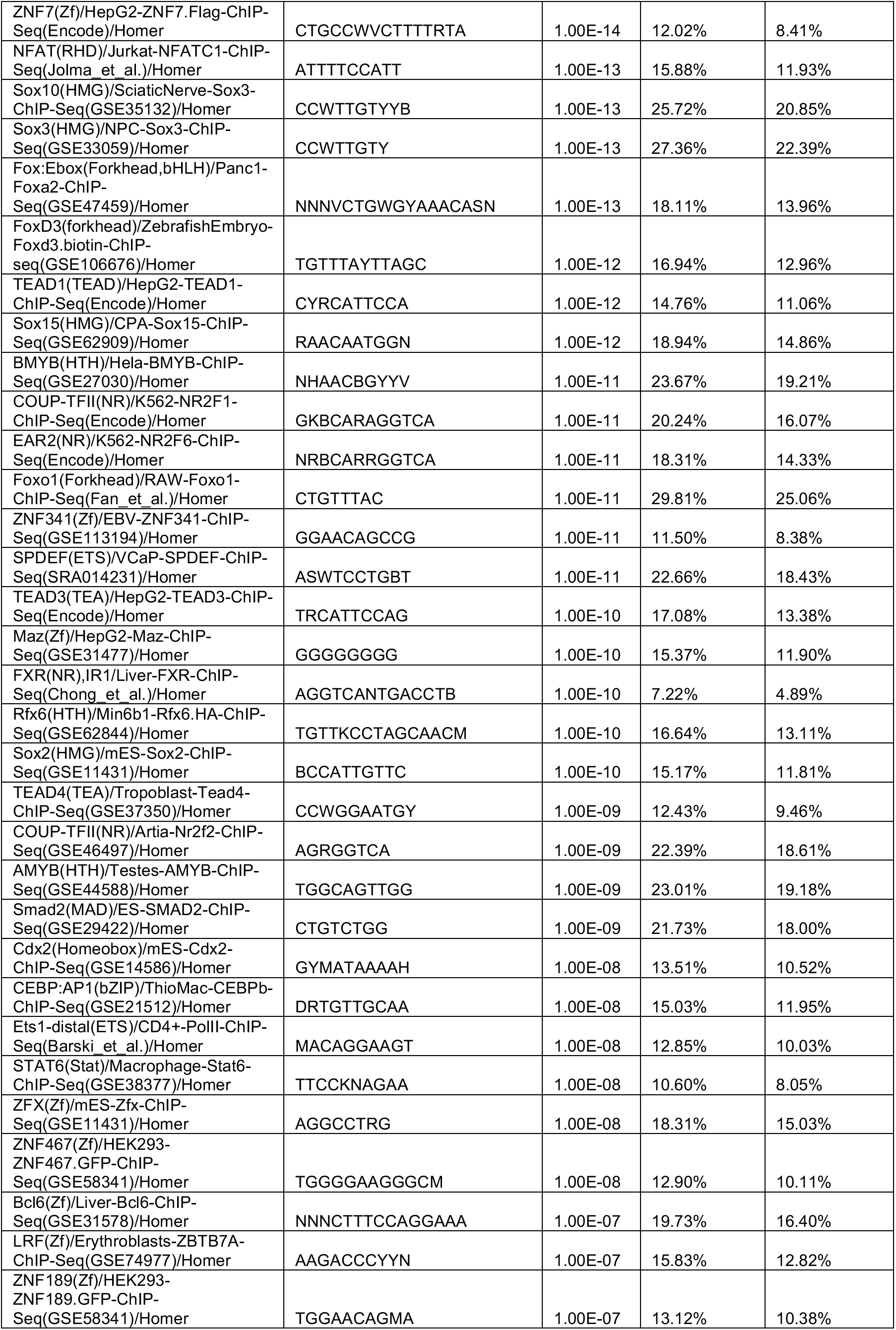
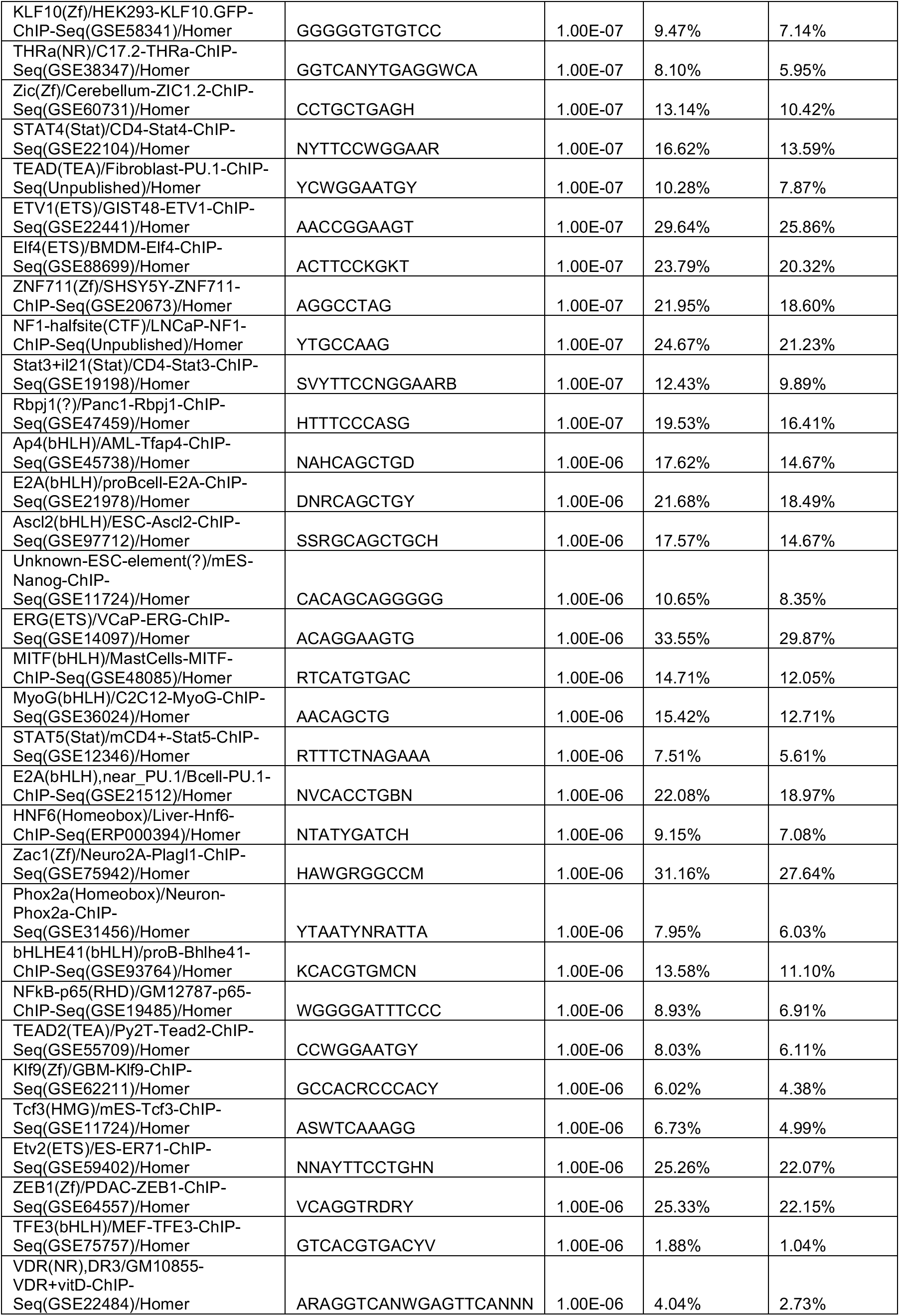
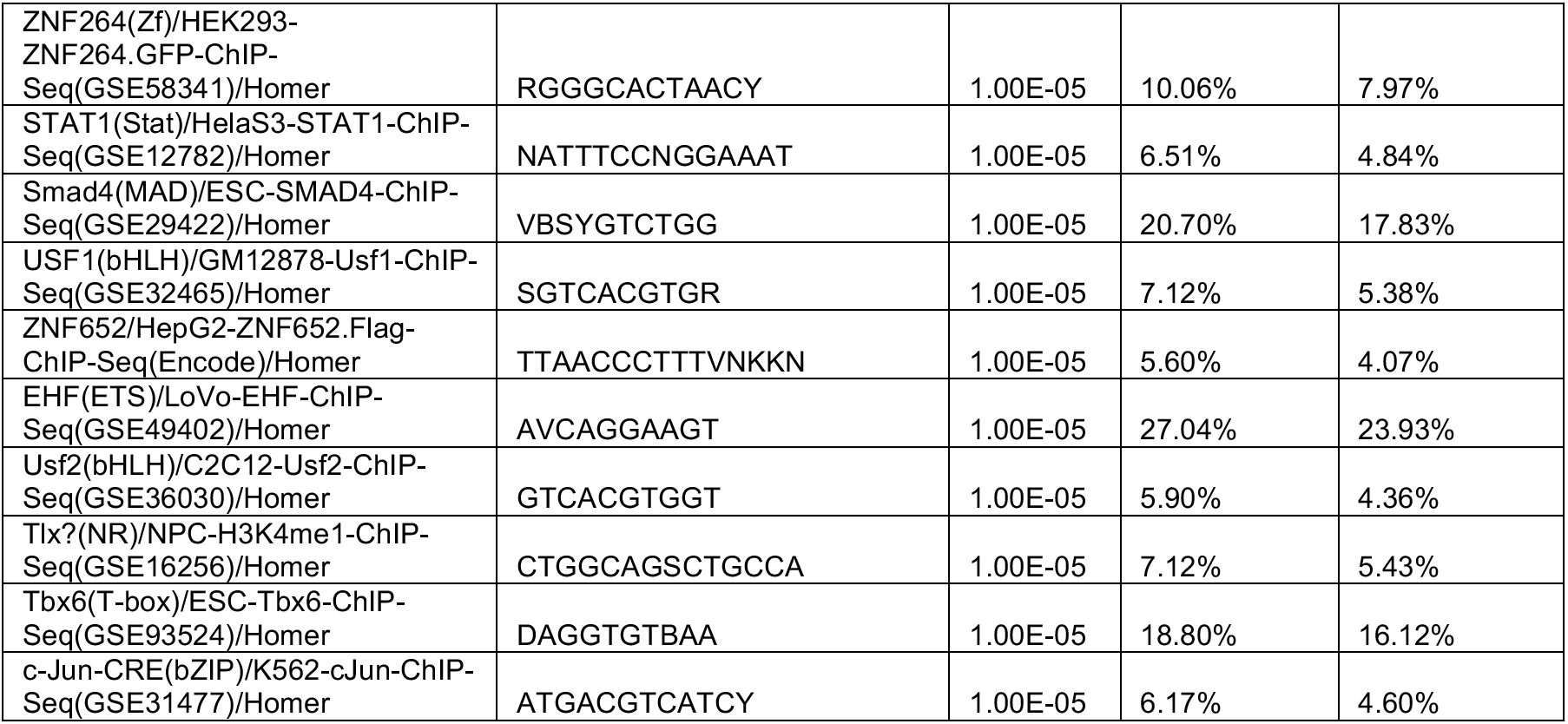
TF motif enrichment in latently infected cells. HOMER TF motif enrichment analysis of differentially open peaks between actively infected cells (GFP+) and latently infected (GFP-) cells. Top 100 enriched motifs in peaks preferentially open in latently infected cells are shown. Target Sequences represent genomic sequences that have significantly elevated accessibility in latently infected cells. Background Sequences represent all open chromatin regions in CD4 T cells.

**Fig S2:**
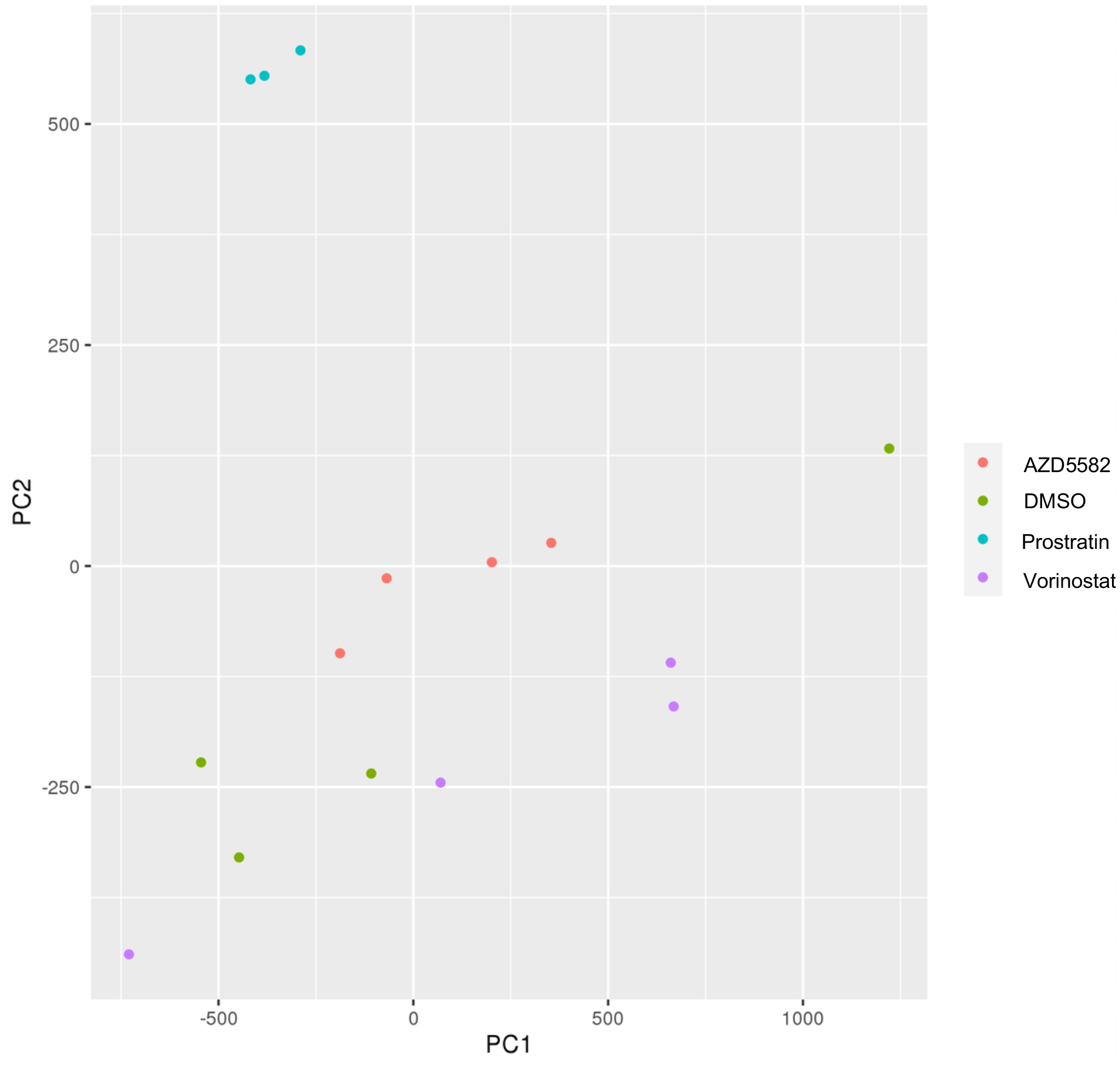
Principal component analysis of infected cells after LRA stimulation ATACseq data. Principal component analysis was performed on ATACseq data from HIV-infected cells after 24h stimulation with three different LRAs or vehicle (DMSO). For all conditions, four independent replicate experiments are shown except for Prostratin, for which three replicates were used. For all LRAs 250nM concentration was used.

**Table S3.**
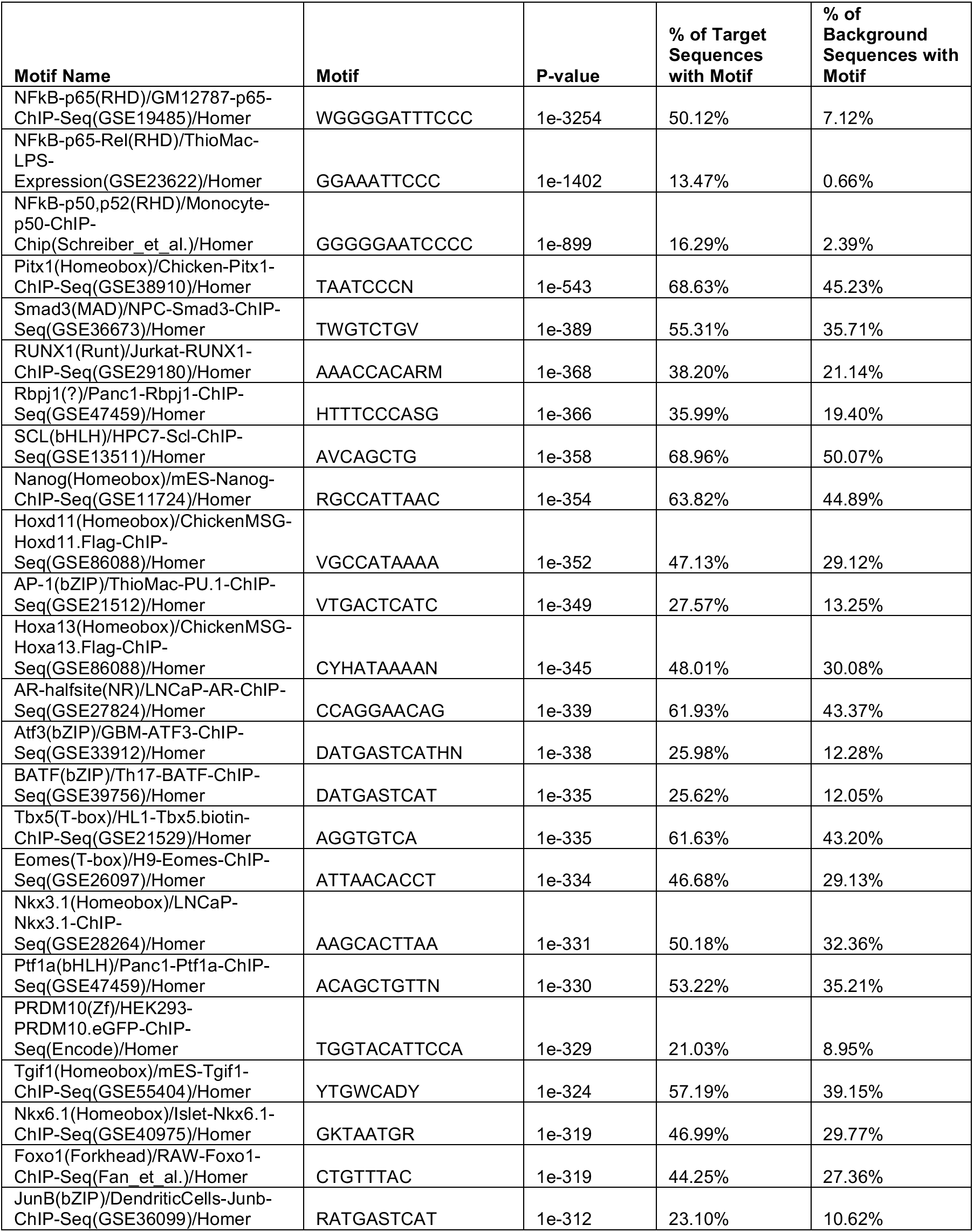
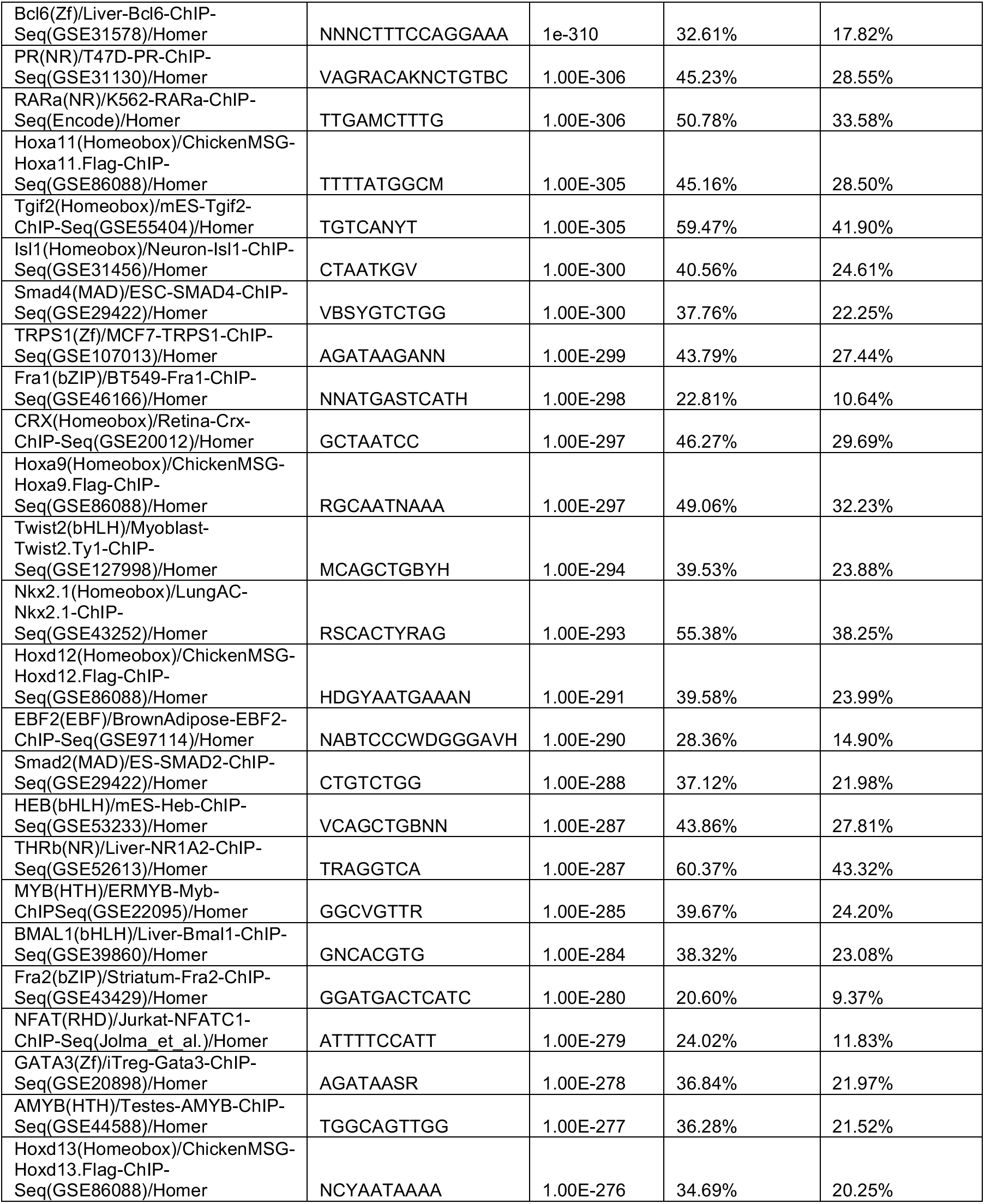
HOMER motif analysis of AZD5582 stimulated latently infected CD4 T cells. The top 50 most highly enriched TF motifs in the set of chromatin peaks that are more open after 24h AZD5582 (250nM) stimulation are shown. Target sequences represent significantly (FDR <0.1) more open chromatin regions after AZD5582 stimulation. Background sequences represent all open chromatin regions in CD4 T cells.

**Table S4.**
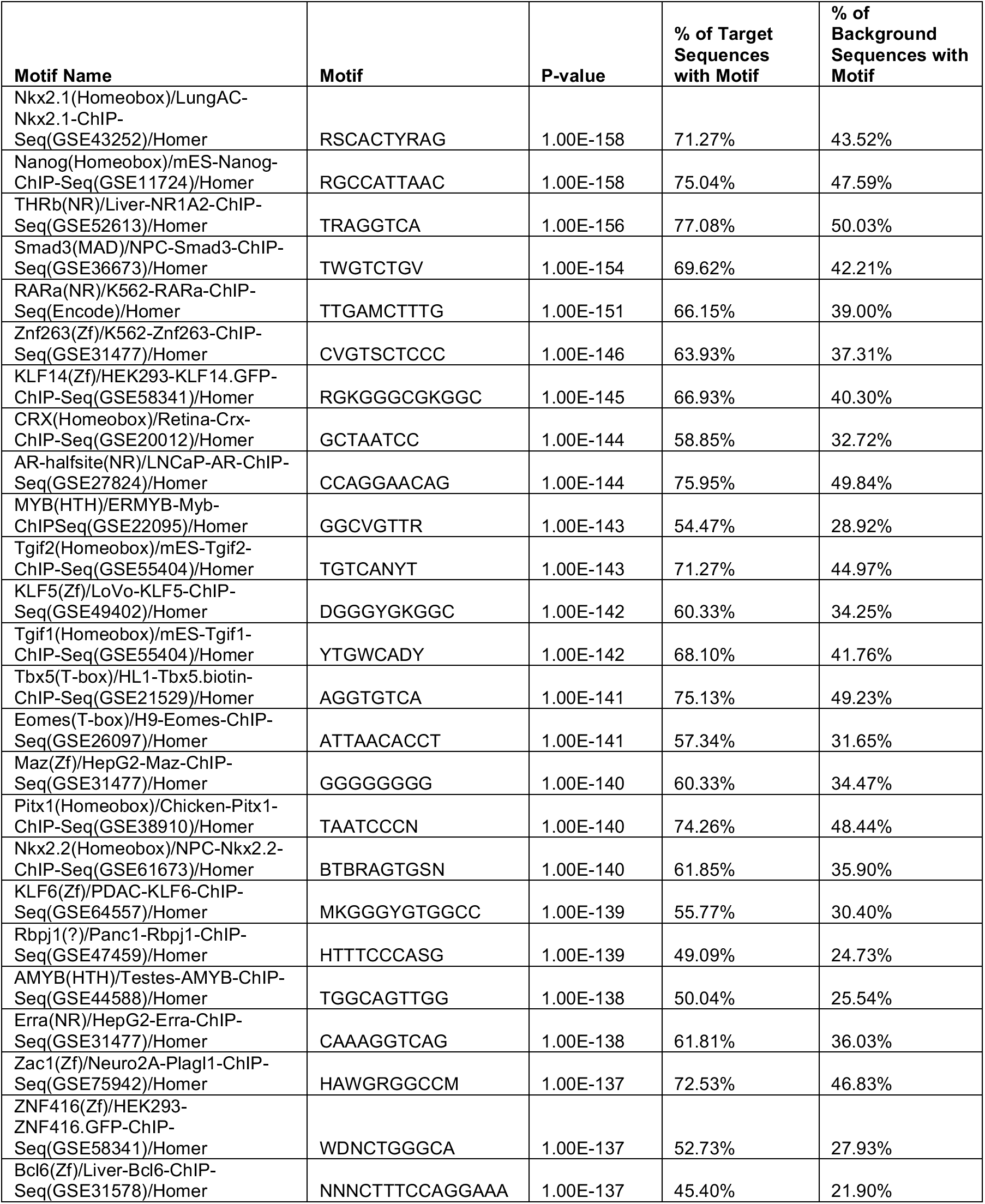
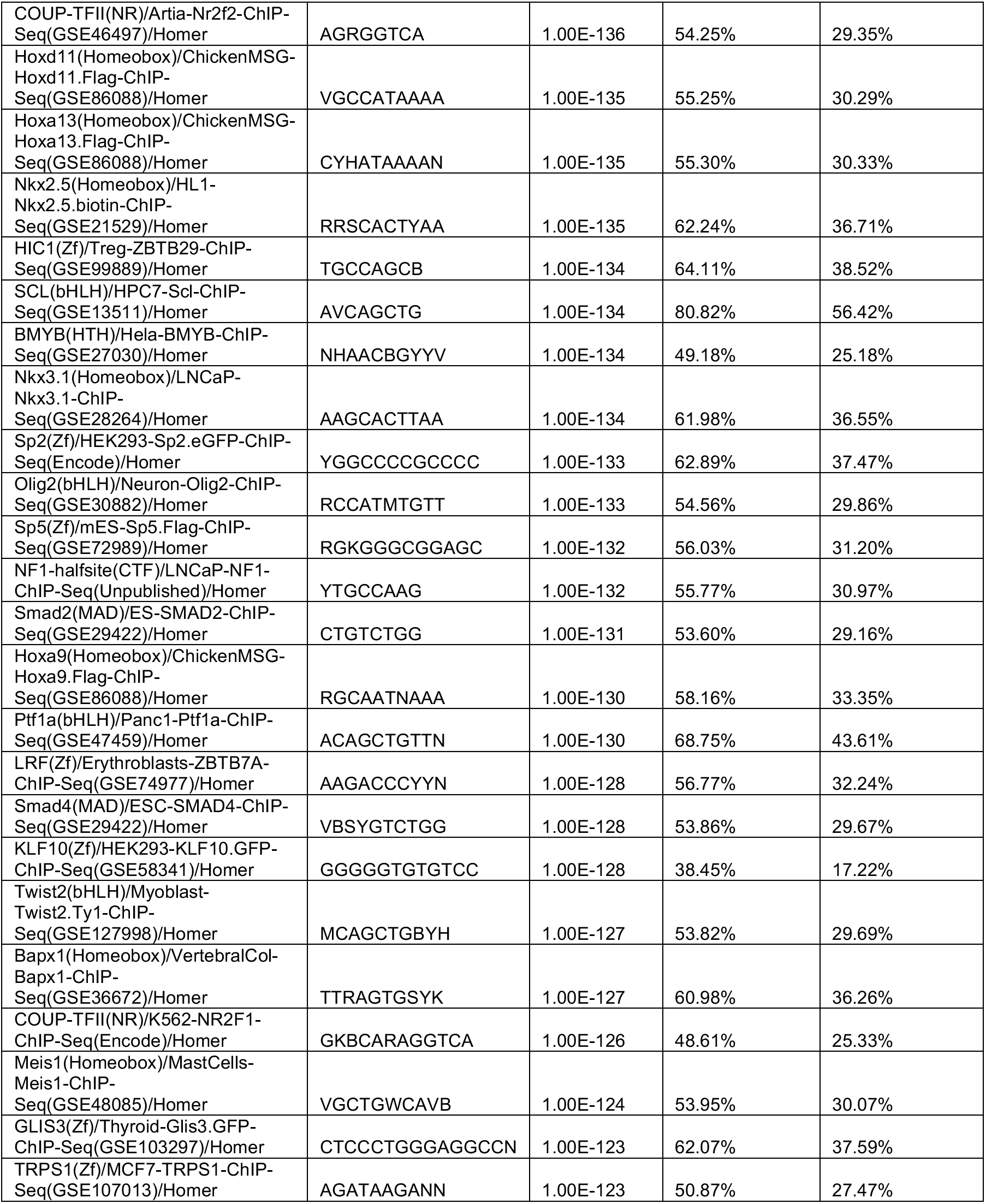
HOMER motif analysis of Vorinostat stimulated latently infected CD4 T cells. The top 50 most highly enriched TF motifs in the set of chromatin peaks that are more open after 24h Vorinostat (250nM) stimulation are shown. Target sequences represent significantly (FDR <0.1) more open chromatin regions after Vorinostat stimulation. Background sequences represent all open chromatin regions in CD4 T cells.

**Table S5.**
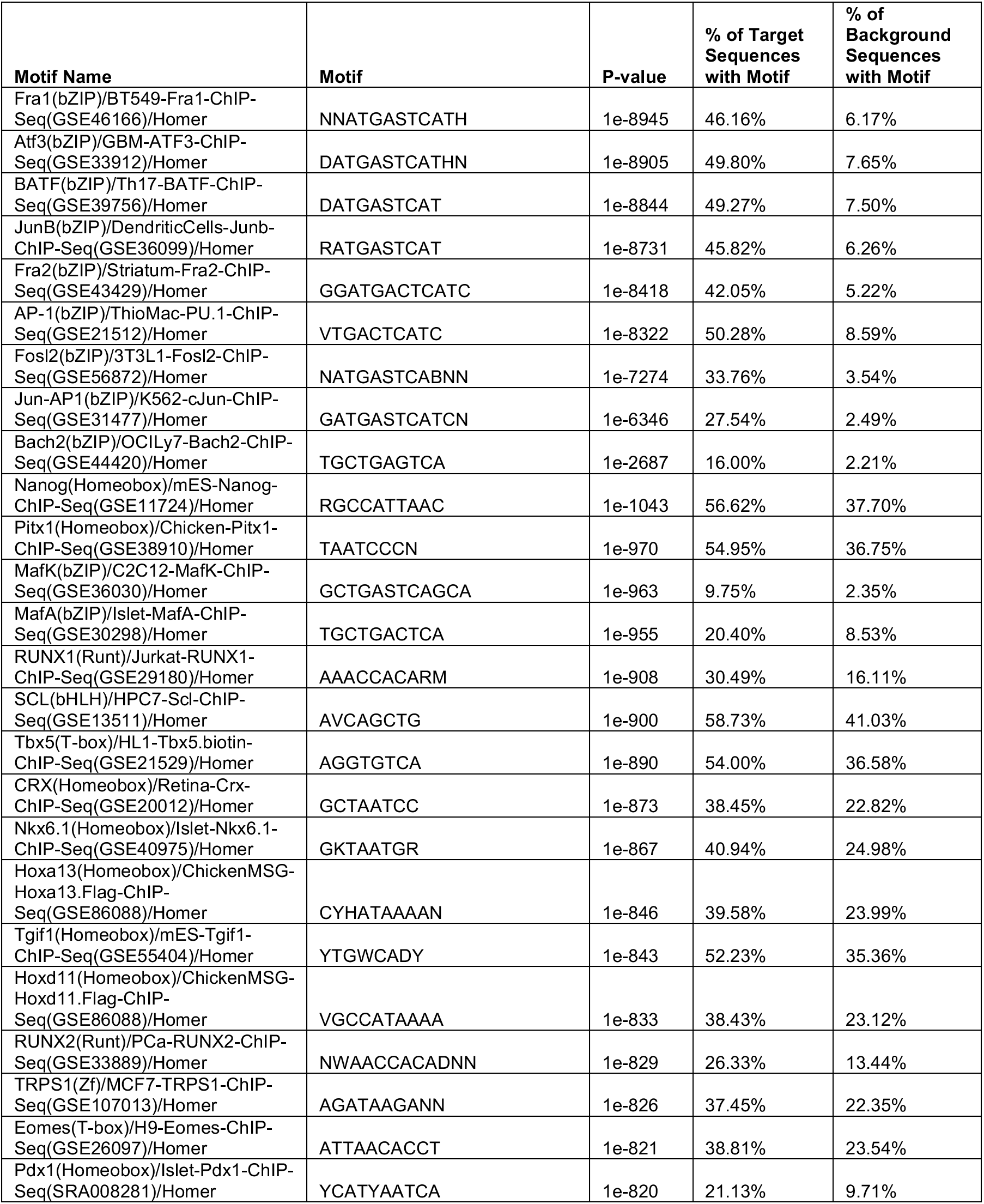
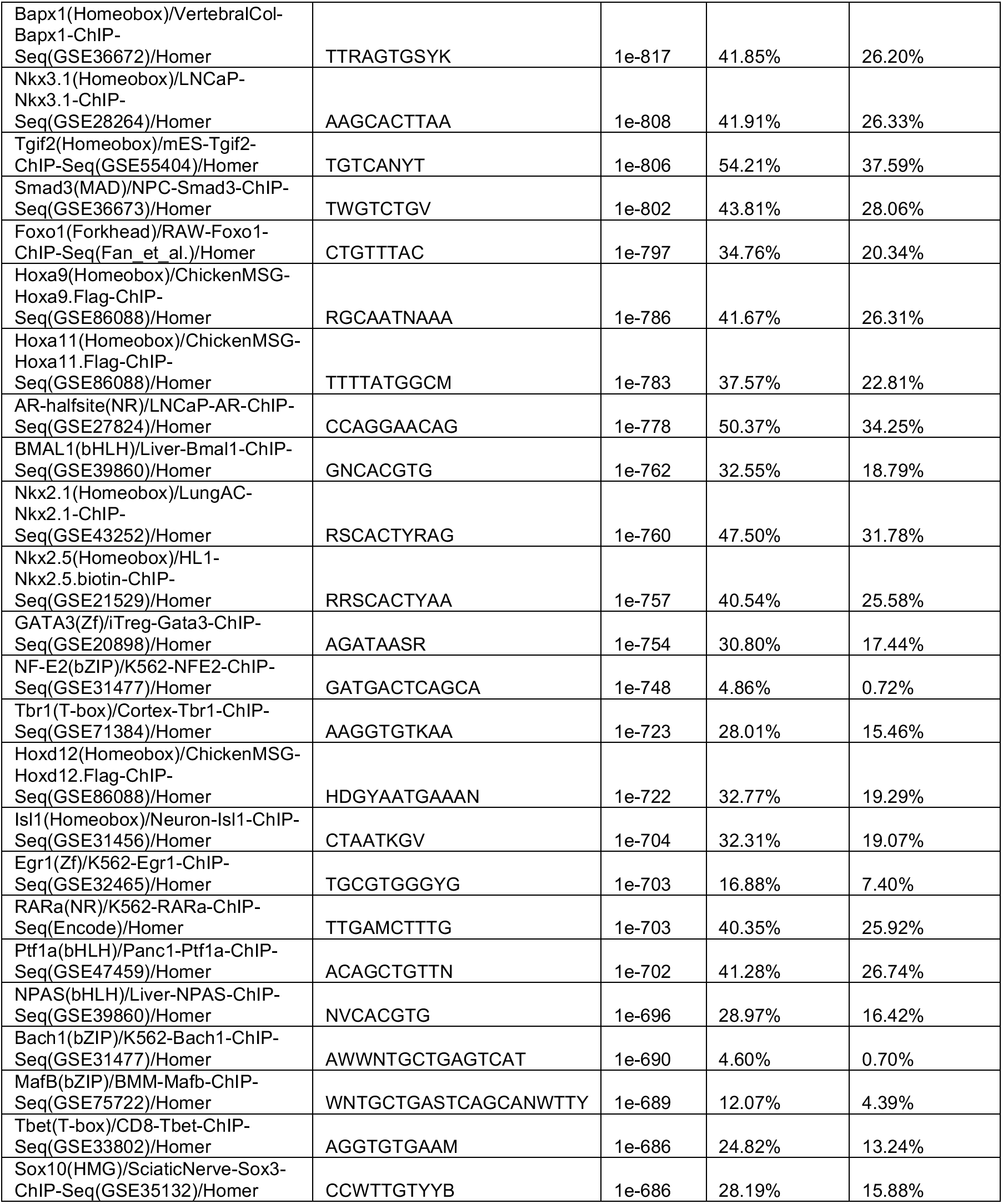
HOMER motif analysis of prostratin stimulated latently infected CD4 T cells. The top 50 most highly enriched TF motifs in the set of chromatin peaks that are more open after 24h prostratin stimulation (250nM) are shown. Target sequences represent significantly (FDR <0.1) more open chromatin regions after prostratin stimulation. Background sequences represent all open chromatin regions in CD4 T cells.

## References

Archin, N., Liberty, A., Kashuba, A., Choudhary, S., Kuruc, J., Crooks, A., Parker, D., Anderson, E., Kearney, M., Strain, M., et al. (2012). Administration of vorinostat disrupts HIV-1 latency in patients on antiretroviral therapy. Nature 487, 482–485.

Barton, K.M., Archin, N.M., Keedy, K.S., Espeseth, A.S., Zhang, Y., Gale, J., Wagner, F.F., Holson, E.B., and Margolis, D.M. (2014). Selective HDAC Inhibition for the Disruption of Latent HIV-1 Infection. PLoS ONE 9.

Battivelli, E., Dahabieh, M.S., Abdel-Mohsen, M., Svensson, J.P., Silva, I.T.D., Cohn, L.B., Gramatica, A., Deeks, S., Greene, W.C., Pillai, S.K., et al. (2018). Chromatin Functional States Correlate with HIV Latency Reversal in Infected Primary CD4+ T Cells. BioRxiv 242958.

Baxter, A.E., Niessl, J., Fromentin, R., Richard, J., Porichis, F., Charlebois, R., Massanella, M., Brassard, N., Alsahafi, N., Delgado, G.-G., et al. (2016). Single-Cell Characterization of Viral Translation-Competent Reservoirs in HIV-Infected Individuals. Cell Host Microbe.

Boeva, V. (2016). Analysis of Genomic Sequence Motifs for Deciphering Transcription Factor Binding and Transcriptional Regulation in Eukaryotic Cells. Front. Genet. 7.

Bradley, T., Ferrari, G., Haynes, B.F., Margolis, D.M., and Browne, E.P. (2018). SingleCell Analysis of Quiescent HIV Infection Reveals Host Transcriptional Profiles that Regulate Proviral Latency. Cell Rep. 25, 107–117.e3.

Buenrostro, J.D., Wu, B., Chang, H.Y., and Greenleaf, W.J. (2015a). ATAC-seq: A Method for Assaying Chromatin Accessibility Genome-Wide. Curr. Protoc. Mol. Biol. 109, 21.29.1–9.

Buenrostro, J.D., Wu, B., Litzenburger, U.M., Ruff, D., Gonzales, M.L., Snyder, M.P., Chang, H.Y., and Greenleaf, W.J. (2015b). Single-cell chromatin accessibility reveals principles of regulatory variation. Nature 523, 486–490.

Chang, L.-H., Ghosh, S., and Noordermeer, D. (2020). TADs and Their Borders: Free Movement or Building a Wall? J. Mol. Biol. 432, 643–652.

Chau, C.M., Zhang, X.-Y., McMahon, S.B., and Lieberman, P.M. (2006). Regulation of Epstein-Barr Virus Latency Type by the Chromatin Boundary Factor CTCF. J. Virol. 80, 5723–5732.

Chomont, N., El-Far, M., Ancuta, P., Trautmann, L., Procopio, F.A., Yassine-Diab, B., Boucher, G., Boulassel, M.-R., Ghattas, G., Brenchley, J.M., et al. (2009). HIV reservoir size and persistence are driven by T cell survival and homeostatic proliferation. Nat. Med. 15, 893–900.

Chougui, G., and Margottin-Goguet, F. (2019). HUSH, a Link Between Intrinsic Immunity and HIV Latency. Front. Microbiol. 10.

Chun, T.-W., Stuyver, L., Mizell, S.B., Ehler, L.A., Mican, J.A.M., Baseler, M., Lloyd, A.L., Nowak, M.A., and Fauci, A.S. (1997). Presence of an inducible HIV-1 latent reservoir during highly active antiretroviral therapy. Proc. Natl. Acad. Sci. 94, 13193–13197.

Crooks, A.M., Bateson, R., Cope, A.B., Dahl, N.P., Griggs, M.K., Kuruc, J.D., Gay, C.L., Eron, J.J., Margolis, D.M., Bosch, R.J., et al. (2015). Precise Quantitation of the Latent HIV-1 Reservoir: Implications for Eradication Strategies. J. Infect. Dis. jiv218.

Dahabieh, M.S., Ooms, M., Brumme, C., Taylor, J., Harrigan, P.R., Simon, V., and Sadowski, I. (2014). Direct non-productive HIV-1 infection in a T-cell line is driven by cellular activation state and NFκB. Retrovirology 11, 17.

Del Rosario, B.C., Kriz, A.J., Del Rosario, A.M., Anselmo, A., Fry, C.J., White, F.M., Sadreyev, R.I., and Lee, J.T. (2019). Exploration of CTCF post-translation modifications uncovers Serine-224 phosphorylation by PLK1 at pericentric regions during the G2/M transition. ELife 8, e42341.

Duverger, A., Wolschendorf, F., Zhang, M., Wagner, F., Hatcher, B., Jones, J., Cron, R.Q., Sluis, R.M. van der, Jeeninga, R.E., Berkhout, B., et al. (2013). An AP-1 Binding Site in \ the Enhancer/Core Element of the HIV-1 Promoter Controls the Ability of HIV-1 To Establish Latent Infection. J. Virol. 87, 2264–2277.

Finzi, D., Hermankova, M., Pierson, T., Carruth, L.M., Buck, C., Chaisson, R.E., Quinn, T.C., Chadwick, K., Margolick, J., Brookmeyer, R., et al. (1997). Identification of a Reservoir for HIV-1 in Patients on Highly Active Antiretroviral Therapy. Science 278, 1295–1300.

Fiorito, E., Sharma, Y., Gilfillan, S., Wang, S., Singh, S.K., Satheesh, S.V., Katika, M.R., Urbanucci, A., Thiede, B., Mills, I.G., et al. (2016). CTCF modulates Estrogen Receptor function through specific chromatin and nuclear matrix interactions. Nucleic Acids Res. 44, 10588–10602.

Friedman, J., Cho, W.-K., Chu, C.K., Keedy, K.S., Archin, N.M., Margolis, D.M., and Karn, J. (2011). Epigenetic Silencing of HIV-1 by the Histone H3 Lysine 27 Methyltransferase Enhancer of Zeste 2▿. J. Virol. 85, 9078–9089.

Garaud, S., Roufosse, F., De Silva, P., Gu-Trantien, C., Lodewyckx, J.-N., Duvillier, H., Dedeurwaerder, S., Bizet, M., Defrance, M., Fuks, F., et al. (2017). FOXP1 is a regulator of quiescence in healthy human CD4+ T cells and is constitutively repressed in T cells from patients with lymphoproliferative disorders. Eur. J. Immunol. 47, 168–179.

Ghirlando, R., and Felsenfeld, G. (2016). CTCF: making the right connections. Genes Dev. 30, 881–891.

Hansen, A.S., Hsieh, T.-H.S., Cattoglio, C., Pustova, I., Saldaña-Meyer, R., Reinberg, D., Darzacq, X., and Tjian, R. (2019). Distinct Classes of Chromatin Loops Revealed by Deletion of an RNA-Binding Region in CTCF. Mol. Cell 76, 395–411.e13.

He, G., Ylisastigui, L., and Margolis, D.M. (2002). The regulation of HIV-1 gene expression: the emerging role of chromatin. DNA Cell Biol. 21, 697–705.

Heinz, S., Benner, C., Spann, N., Bertolino, E., Lin, Y.C., Laslo, P., Cheng, J.X., Murre, C., Singh, H., and Glass, C.K. (2010). Simple combinations of lineage-determining transcription factors prime cis-regulatory elements required for macrophage and B cell identities. Mol. Cell 38, 576–589.

Holwerda, S.J.B., and de Laat, W. (2013). CTCF: the protein, the binding partners, the binding sites and their chromatin loops. Philos. Trans. R. Soc. B Biol. Sci. 368.

Kim, Y.K., Mbonye, U., Hokello, J., and Karn, J. (2011). T-cell receptor signaling enhances transcriptional elongation from latent HIV proviruses by activating P-TEFb through an ERK-dependent pathway. J. Mol. Biol. 410, 896–916.

Lederman, M.M., Calabrese, L., Funderburg, N.T., Clagett, B., Medvik, K., Bonilla, H., Gripshover, B., Salata, R.A., Taege, A., Lisgaris, M., et al. (2011). Immunologic Failure Despite Suppressive Antiretroviral Therapy Is Related to Activation and Turnover of Memory CD4 Cells. J. Infect. Dis. 204, 1217–1226.

Lederman, M.M., Funderburg, N.T., Sekaly, R.P., Klatt, N.R., and Hunt, P.W. (2013). Residual Immune Dysregulation Syndrome in Treated HIV infection. Adv. Immunol. 119, 51–83.

Lee, G.Q., and Lichterfeld, M. (2016). Diversity of HIV-1 reservoirs in CD4 T cell subsets. Curr. Opin. HIV AIDS 11, 383–387.

Li, H. (2013). Aligning sequence reads, clone sequences and assembly contigs with BWA-MEM. ArXiv13033997 Q-Bio.

Li, H., Handsaker, B., Wysoker, A., Fennell, T., Ruan, J., Homer, N., Marth, G., Abecasis, G., Durbin, R., and 1000 Genome Project Data Processing Subgroup (2009). The Sequence Alignment/Map format and SAMtools. Bioinforma. Oxf. Engl. 25, 2078–2079.

Lichtfuss, G.F., Hoy, J., Rajasuriar, R., Kramski, M., Crowe, S.M., and Lewin, S.R. (2011). Biomarkers of immune dysfunction following combination antiretroviral therapy for HIV infection. Biomark. Med. 5, 171–186.

Lun, A.T.L., and Smyth, G.K. (2016). csaw: a Bioconductor package for differential binding analysis of ChIP-seq data using sliding windows. Nucleic Acids Res. 44, e45.

Maldarelli, F., Wu, X., Su, L., Simonetti, F.R., Shao, W., Hill, S., Spindler, J., Ferris, A.L., Mellors, J.W., Kearney, M.F., et al. (2014). Specific HIV integration sites are linked to clonal expansion and persistence of infected cells. Science 345, 179–183.

Margolis, D.M. (2014). How Might We Cure HIV? Curr. Infect. Dis. Rep. 16, 392.

McCarthy, D.J., Chen, Y., and Smyth, G.K. (2012). Differential expression analysis of multifactor RNA-Seq experiments with respect to biological variation. Nucleic Acids Res. 40, 4288–4297.

Mousseau, G., and Valente, S.T. (2015). Didehydro-Cortistatin A: a new player in HIV-therapy? Expert Rev. Anti Infect. Ther. 0, null.

Mzingwane, M.L., and Tiemessen, C.T. (2017). Mechanisms of HIV persistence in HIV reservoirs. Rev. Med. Virol. 27.

Nabel, G., and Baltimore, D. (1987). An inducible transcription factor activates expression of human immunodeficiency virus in T cells. Nature 326, 711–713.

Nixon, C.C., Mavigner, M., Sampey, G.C., Brooks, A.D., Spagnuolo, R.A., Irlbeck, D.M., Mattingly, C., Ho, P.T., Schoof, N., Cammon, C.G., et al. (2020). Systemic HIV and SIV latency reversal via non-canonical NF-κB signalling in vivo. Nature 578, 160–165.

Oh, S.A., Seki, A., and Rutz, S. (2019). Ribonucleoprotein Transfection for CRISPR/Cas9-Mediated Gene Knockout in Primary T Cells. Curr. Protoc. Immunol. 124, e69.

Pearson, R., Kim, Y.K., Hokello, J., Lassen, K., Friedman, J., Tyagi, M., and Karn, J. (2008). Epigenetic Silencing of Human Immunodeficiency Virus (HIV) Transcription by Formation of Restrictive Chromatin Structures at the Viral Long Terminal Repeat Drives the Progressive Entry of HIV into Latency. J. Virol. 82, 12291–12303.

Pinkevych, M., Cromer, D., Tolstrup, M., Grimm, A.J., Cooper, D.A., Lewin, S.R., Søgaard, O.S., Rasmussen, T.A., Kent, S.J., Kelleher, A.D., et al. (2015). HIV Reactivation from Latency after Treatment Interruption Occurs on Average Every 5-8 Days—Implications for HIV Remission. PLoS Pathog. 11.

Rafati, H., Parra, M., Hakre, S., Moshkin, Y., Verdin, E., and Mahmoudi, T. (2011). Repressive LTR Nucleosome Positioning by the BAF Complex Is Required for HIV Latency. PLoS Biol. 9.

Razooky, B.S., Pai, A., Aull, K., Rouzine, I.M., and Weinberger, L.S. (2015). A Hardwired HIV Latency Program. Cell 160, 990–1001.

Reeves, D.B., Duke, E.R., Wagner, T.A., Palmer, S.E., Spivak, A.M., and Schiffer, J.T. (2018). A majority of HIV persistence during antiretroviral therapy is due to infected cell proliferation. Nat. Commun. 9.

Roebuck, K.A., Gu, D.S., and Kagnoff, M.F. (1996). Activating protein-1 cooperates with phorbol ester activation signals to increase HIV-1 expression. AIDS Lond. Engl. 10, 819–826.

Roux, A., Leroy, H., Muylder, B.D., Bracq, L., Oussous, S., Dusanter-Fourt, I., Chougui, G., Tacine, R., Randriamampita, C., Desjardins, D., et al. (2019). FOXO1 transcription factor plays a key role in T cell—HIV-1 interaction. PLOS Pathog. 15, e1007669.

Saldaña-Meyer, R., Rodriguez-Hernaez, J., Escobar, T., Nishana, M., Jácome-López, K., Nora, E.P., Bruneau, B.G., Tsirigos, A., Furlan-Magaril, M., Skok, J., et al. (2019). RNA Interactions Are Essential for CTCF-Mediated Genome Organization. Mol. Cell 76, 412–422.e5.

Seki, A., and Rutz, S. (2018). Optimized RNP transfection for highly efficient CRISPR/Cas9-mediated gene knockout in primary T cells. J. Exp. Med. 215, 985–997.

Serrano-Villar, S., Sainz, T., Lee, S.A., Hunt, P.W., Sinclair, E., Shacklett, B.L., Ferre, A.L., Hayes, T.L., Somsouk, M., Hsue, P.Y., et al. (2014). HIV-Infected Individuals with Low CD4/CD8 Ratio despite Effective Antiretroviral Therapy Exhibit Altered T Cell Subsets, Heightened CD8+ T Cell Activation, and Increased Risk of Non-AIDS Morbidity and Mortality. PLoS Pathog. 10.

Shan, L., Xing, S., Yang, H.-C., Zhang, H., Margolick, J.B., and Siliciano, R.F. (2014). Unique characteristics of histone deacetylase inhibitors in reactivation of latent HIV-1 in Bcl-2-transduced primary resting CD4+ T cells. J. Antimicrob. Chemother. 69, 28–33.

Siliciano, R.F., and Greene, W.C. (2011). HIV Latency. Cold Spring Harb. Perspect. Med. 1, a007096.

Siliciano, J.D., Kajdas, J., Finzi, D., Quinn, T.C., Chadwick, K., Margolick, J.B., Kovacs, C., Gange, S.J., and Siliciano, R.F. (2003). Long-term follow-up studies confirm the stability of the latent reservoir for HIV-1 in resting CD4+ T cells. Nat. Med. 9, 727–728.

Søgaard, O.S., Graversen, M.E., Leth, S., Olesen, R., Brinkmann, C.R., Nissen, S.K., Kjaer, A.S., Schleimann, M.H., Denton, P.W., Hey-Cunningham, W.J., et al. (2015). The Depsipeptide Romidepsin Reverses HIV-1 Latency In Vivo. PLoS Pathog 11, e1005142.

Soriano-Sarabia, N., Bateson, R.E., Dahl, N.P., Crooks, A.M., Kuruc, J.D., Margolis, D.M., and Archin, N.M. (2014). Quantitation of Replication-Competent HIV-1 in Populations of Resting CD4+ T Cells. J. Virol. 88, 14070–14077.

Strahl, B.D., and Allis, C.D. (2000). The language of covalent histone modifications. Nature 403, 41–45.

Thorvaldsdóttir, H., Robinson, J.T., and Mesirov, J.P. (2013). Integrative Genomics Viewer (IGV): high-performance genomics data visualization and exploration. Brief. Bioinform. 14, 178–192.

Tripathy, M.K., McManamy, M.E.M., Burch, B.D., Archin, N.M., and Margolis, D.M. (2015). H3K27 Demethylation at the Proviral Promoter Sensitizes Latent HIV to the Effects of Vorinostat in Ex Vivo Cultures of Resting CD4+ T Cells. J. Virol. 89, 8392–8405.

Tyagi, M., Pearson, R.J., and Karn, J. (2010). Establishment of HIV Latency in Primary CD4+ Cells Is due to Epigenetic Transcriptional Silencing and P-TEFb Restriction. J. Virol. 84, 6425–6437.

Vallejo-Gracia, A., Chen, I.P., Perrone, R., Besnard, E., Boehm, D., Battivelli, E., Tezil, T., Krey, K., Raymond, K.A., Hull, P.A., et al. (2020). FOXO1 promotes HIV latency by suppressing ER stress in T cells. Nat. Microbiol.

Wang, H., Maurano, M.T., Qu, H., Varley, K.E., Gertz, J., Pauli, F., Lee, K., Canfield, T., Weaver, M., Sandstrom, R., et al. (2012). Widespread plasticity in CTCF occupancy linked to DNA methylation. Genome Res. 22, 1680–1688.

Washington, S.D., Edenfield, S.I., Lieux, C., Watson, Z.L., Taasan, S.M., Dhummakupt, A., Bloom, D.C., and Neumann, D.M. (2018). Depletion of the Insulator Protein CTCF Results in Herpes Simplex Virus 1 Reactivation In Vivo. J. Virol. 92.

Weinberger, L.S., Burnett, J.C., Toettcher, J.E., Arkin, A.P., and Schaffer, D.V. (2005). Stochastic Gene Expression in a Lentiviral Positive-Feedback Loop: HIV-1 Tat Fluctuations Drive Phenotypic Diversity. Cell 122, 169–182.

Yang, Z., and Engel, J.D. (1993). Human T cell transcription factor GATA-3 stimulates HIV-1 expression. Nucleic Acids Res. 21, 2831–2836.

Yang, H.-C., Xing, S., Shan, L., O’Connell, K., Dinoso, J., Shen, A., Zhou, Y., Shrum, C.K., Han, Y., Liu, J.O., et al. (2009). Small-molecule screening using a human primary cell model of HIV latency identifies compounds that reverse latency without cellular activation. J. Clin. Invest. 119, 3473–3486.

Yang, X., Chen, Y., and Gabuzda, D. (1999). ERK MAP kinase links cytokine signals to activation of latent HIV-1 infection by stimulating a cooperative interaction of AP-1 and NF-kappaB. J. Biol. Chem. 274, 27981–27988.

Yoshimura, K. (2017). Current status of HIV/AIDS in the ART era. J. Infect. Chemother. Off. J. Jpn. Soc. Chemother. 23, 12–16.

